# Recording proteome dynamics in the late-stage embryo at hour-timescale, spatial, and single-cell resolution with MEMBRYO

**DOI:** 10.64898/2026.09.21.753234

**Authors:** Marta Couce-Iglesias, Eleonora Oliani, Robert Kerridge, Florian Mutschler, Daniela S. Valdes, Anna Sophie Welter, Beata Lukaszewska-McGreal, David Meierhofer, Koshi Imami, Eugenio F. Fornasiero, Fabian Coscia, Matthias Selbach, Matthew L. Kraushar

## Abstract

During embryonic development, cells undergo extensive proteome remodeling as they acquire specialized identities and assemble into complex tissues. Yet how the selectivity of protein synthesis, degradation, and inheritance from earlier cellular states collectively shape cell identity remains largely unknown, owing to the difficulty of tracking proteome dynamics within developing embryos. Here we introduce MEMBRYO, an embryo culture platform enabling fast and efficient metabolic labeling of newly synthesized proteins throughout live mouse embryos. By extending embryo culture to previously inaccessible stages approaching birth, MEMBRYO enables the investigation of proteome dynamics from early to advanced stages of development, including differentiation of the cerebral cortex. Combining whole-embryo stable isotope labeling with quantitative proteomics, we uncover widespread protein synthesis and degradation dynamics in the developing cortex that are uncoupled from transcriptional changes. Deep labeling further enables time-resolved, spatial, and single-cell proteomics, distinguishing newly synthesized from pre-existing proteins in diverse cells. Although Ncam1 and Plch1 both accumulate during the differentiation of neural progenitors, their protein pools exhibit opposing renewal dynamics. Pre-existing Ncam1 is increasingly retained during differentiation, consistent with the selective stabilization of membrane adhesion proteins, whereas the calcium-signaling enzyme Plch1 is increasingly renewed. Together, our findings reveal how neuronal identity emerges through a balance of proteome plasticity and stability, establishing MEMBRYO as a versatile platform for investigating the dynamic molecular basis of embryonic development.

## Introduction

The embryo is formed by dynamic molecular states that drive cellular phenotypes. The maintenance of progenitor states, or the transition towards new differentiated states, are enacted by the synthesis of new proteins and degradation of old ones ^1^. As such, protein synthesis and degradation, which combined represent protein turnover kinetics ^2–4^, point toward the direction a cell will take in development. Tracking proteome kinetics over time is a major challenge in the complex cellular environment of the embryo, especially at the hour-timescales of developmental phenotypes. Since conventional proteomics measures a snapshot of protein abundance at the time of collection, most current molecular models of embryonic development lack kinetic, time-continuous, proteome activity *in vivo*.

Technologies that gain kinetic molecular information require direct access to the cell to label its molecules ^4,5^. In these methods, cells are pulsed with analogs of nucleic or amino acids to label newly synthesized mRNA or protein, respectively, which encode a detectable signature. Nascent protein synthesis and turnover can be tracked with “heavy” stable isotopic amino acids for detection by mass spectrometry (SILAC-MS) ^4,5^. Proteome synthesis and turnover are calculated by direct comparison of pulse labeled (new) vs. unlabeled (old) protein in the same sample. Stable isotope labeling techniques can be used to study long-term protein turnover in animal models including mice ^2,6^, but these technologies have had limited applications in the embryo or tissues like the developing brain. Direct access of the analogs to cells in the embryo *in utero* is a major obstacle, leading to low (∼10%) proteome labeling in a prior attempt ^7^. Consequently, our concept of gene expression kinetics is mainly derived from cells in culture ^4^, and is largely unknown in neurodevelopment ^1^.

Here we advance a mouse embryo culture method for kinetic proteome analysis by SILAC-MS called MEMBRYO (<u>M</u>etabolic labeling in the <u>EMBRYO</u>). The method extends the period of embryo culture into the latest stages of organogenesis, which we employ to analyze proteome kinetics in the last region of the nervous system to develop in mammals – the cortex. Continuous SILAC labeling of the nascent proteome achieves highly comprehensive tracking of protein synthesis and turnover, and uncovered multiple layers of post-transcriptional gene expression regulation in the cortex. Protein synthesis and turnover frequently diverge from patterns of transcription for key milestones of neurodevelopment. MEMBRYO’s exceptional labeling depth extended SILAC proteomics into both spatial and single-cell dimensions *in vivo*, which revealed kinetic protein information in neural progenitors and differentiating neurons. Protein turnover variations affect key neurodevelopmental pathways, including Ncam1 associated with synaptogenesis. Thus, MEMBRYO accesses time-continuous protein kinetics in the embryo and advances our molecular models neurodevelopment.

## Results

The foundation of our method is the *ex utero* culture of mouse embryos (**Fig. 1a**), which has traditionally been limited to developmental stages up to E11.5 ^8–10^. Here, we extend this capability to later embryonic stages E12.5 to E16.5 by refining the roller culture technique ^9,10^. Accessing later developmental stages allows for a comprehensive analysis of organogenesis up to and including neurogenesis in the mouse brain, which occurs throughout the last gestational trimester ^1^.

**Fig. 1.**
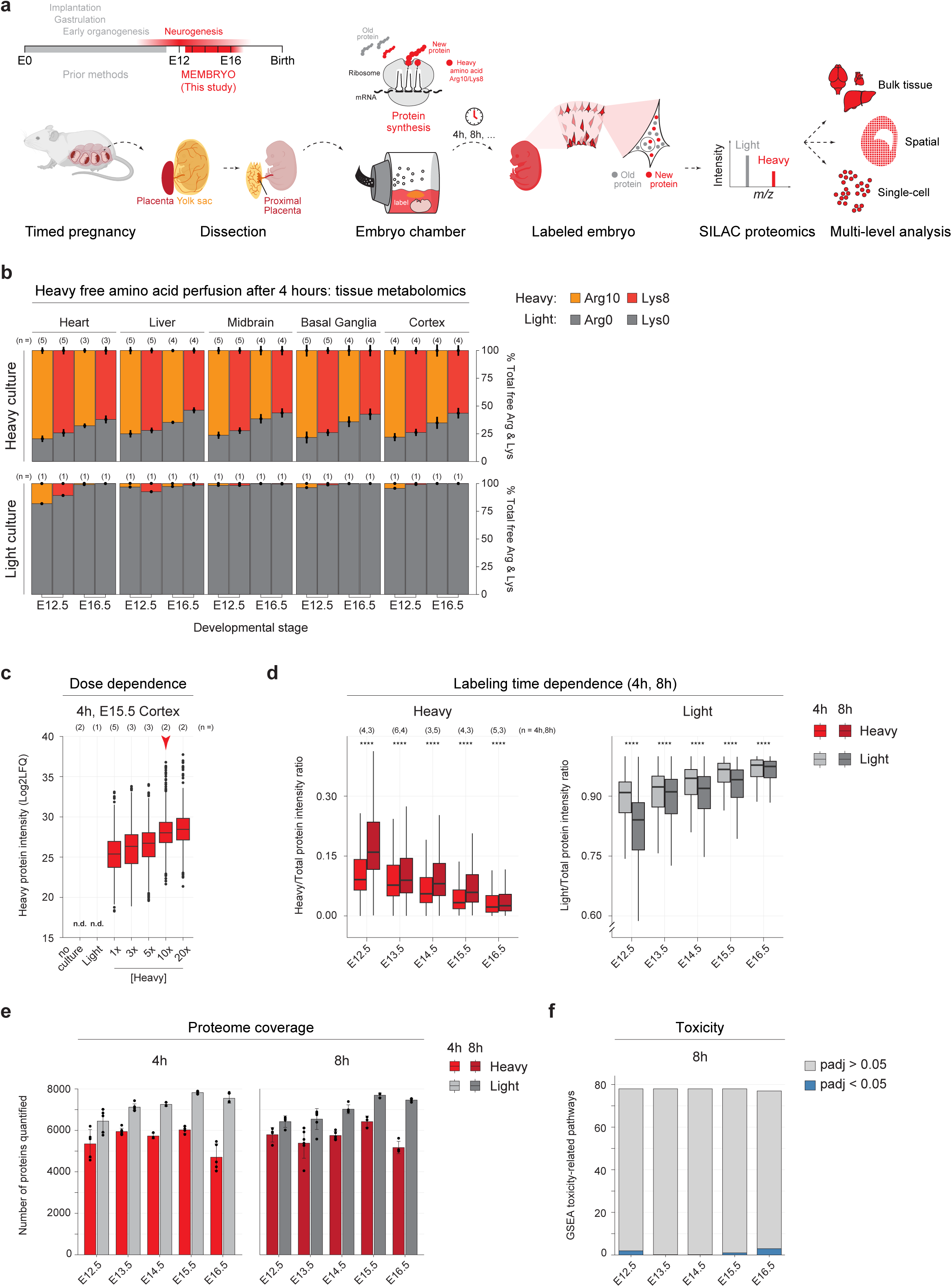
Metabolic labeling of newly synthesized proteins in embryo culture. **a**, Schematic overview of MEMBRYO for SILAC proteomics analysis of the mouse embryo, including the period of neurogenesis in the forebrain. Dissected embryos were cultured *ex utero* in medium supplemented with heavy isotope arginine and lysine (Arg10/Lys8) for labeling newly synthesized proteins, which are quantified by bulk, spatial, and single-cell proteomics in this study. Icons were adapted from BioRender (https://BioRender.com/4p0hww9). **b**, Metabolomics quantification of heavy analog levels in five embryonic tissues (heart, liver, midbrain, basal ganglia, and cortex) after 4 hours of culture at E12.5 and E16.5. Embryos cultured with light analogs (Arg0/Lys0) are negative controls. The percent of total arginine and lysine that is heavy labeled or the light counterpart is shown, with biological replicate (n) embryos for each mean ± standard deviation. Biological samples were measured in three technical replicates. **c**, SILAC-MS analysis of cortical tissue after 4 h of culture with increasing concentrations of heavy amino acid analogs (1x = 400 µM arginine and 800 µM lysine). Biological replicate (n) embryos are shown. **d**, Heavy labeled protein intensity in cortical tissue measured by SILAC-MS after 4 and 8 hours of culture at each stage from E12.5-E16.5. Light protein intensity corresponds to pre-existing unlabeled protein. Differences between 4 and 8 hours were assessed separately at each embryonic stage with a two-sided unpaired Wilcoxon rank-sum tests. P values were adjusted for multiple comparisons with the Benjamini-Hochberg method. Biological replicate (n) embryos are shown. **e**, Number of proteins quantified by SILAC-MS after 4 or 8 hours of culture from E12.5 to E16.5. Red bars indicate newly synthesized heavy labeled proteins, while grey bars indicate the corresponding unlabeled pool. Points represent individual biological replicates. **f**, Number of toxicity-associated gene sets enriched after 8 h of SILAC labeling, compared to uncultured/unlabeled fresh tissue, at each developmental stage. Blue bars indicate significantly enriched toxicity pathways, while grey bars indicate all tested toxicity-related pathways.

Viable embryo culture at late developmental stages is supported by our dissection strategy, which retains the umbilical cord, its proximal connection to the placenta, and the vascularized yolk sac (**Fig. 1a**). The interface between the umbilical cord and placenta must be maintained – some placental tissue is required to provide a scaffold that keeps vessel ends open. Maintaining the vessel patency allows the embryo’s beating heart to circulate the surrounding media and perfuse its tissues, while preventing cardiovascular collapse. The surrounding media can be reconstituted with a variety of molecules that perfuse the live embryo, such as SILAC amino acids. The culture chamber maintains temperature and gas partial pressure, while agitating (rolling) the mixture to remove metabolic waste and recirculate new molecules. Embryo survival rates after 4-8 hours of culture at stages between E12.5-E16.5 range from 100% ∼ 25% survival on average per litter, and result in fewer successful embryos with longer incubation times and at later embryonic stages (**Extended Data Fig. 1a-b**) – consistent with previous cultures up to 11.5 ^8–10^. Embryos were considered healthy and committed to further analysis only if a clear heartbeat and fluid movement in the umbilical cord was observed at the end of the experiment, with minimal hemorrhage or interstitial fluid accumulation. These conditions were the basis for optimizing proteome-wide SILAC labeling of the embryo at time scales like 4-8 hours that are consistent with protein synthesis rates ^5^ (**Fig. 1a**).

### Heavy amino acid labeling and SILAC proteomics in cultured embryos

We first tested whether embryos cultured for 4 hours are perfused by heavy (Arg10, Lys8) SILAC amino acids into a variety of organs (**Fig. 1b**). Free amino acids (unincorporated into peptides) were measured by metabolomic MS, representing the available amino acid supply for protein synthesis-based labeling. Heavy free amino acids were analyzed in the heart, liver, midbrain, basal ganglia, and cortex, and compared to pre-existing light free amino acids (Arg0, Lys0). The amount of heavy vs. light in each tissue was calculated as a percent of the total. Analysis of tissues at E12.5 and E16.5 showed that heavy replaced ∼50-75% of the light amino acid pool after 4 hours, with consistent levels of incorporation between organs. Embryos cultured with only light amino acids demonstrated minimal heavy SILAC detection as a negative control. The replacement of light with heavy amino acids was ∼ 15% less efficient at E16.5 compared to E12.5 in SILAC cultures. However, lower heavy amino acid levels correlate directly with a decrease in pre-existing light amino acids from E12.5-E16.5 (**Extended Data Fig. 1c-d**). Along with prior work that found progressively decreasing histidine in the embryonic brain ^11^, these data suggest that lower free amino acid availability late in organogenesis is physiologic. Taken together, we demonstrate that MEMBRYO reconstitutes the majority of free amino acids in cultured embryos within 4 hours for acute protein synthesis labeling.

We next optimized the concentration of heavy SILAC in the media to reach peak labeling efficiency of the newly synthesized proteome in the embryo (**Fig. 1c**). E15.5 embryos were labeled with increasing concentrations of heavy SILAC, above the typical range (1x) used for cell lines (400 μM Arg10, 800 μM Lys8) ^12^. MS analysis of cortical tissue demonstrated a concentration-dependent increase in proteome labeling intensity (LFQ), reaching a plateau at 10x concentration. These data provided a minimally effective concentration for all future experiments, and confirm that proteome labeling in the embryo with SILAC is dose-dependent.

We next assessed whether increasing the labeling duration results in increased proteome labeling intensity. 4- and 8-hour cultures at each stage of cortical neurogenesis (E12.5-E16.5) were analyzed for heavy labeling intensity in the cortical tissue (**Fig. 1d**). Heavy labeling increases from 4 to 8 hours at each stage, with a concomitant decrease in light as expected from its turnover. We reliably quantified ∼ 5580 heavy proteins per stage (**Fig. 1e**), demonstrating highly comprehensive proteome labeling at time scales reflecting protein synthesis and turnover. Relatively low arginine-to-proline conversion rates (∼1%) and high labeling efficiency per-residue (∼82%) in cortical tissue across all stages further support the fidelity and depth of embryo labeling (**Extended Data Fig. 1e-f**).

Finally, we assessed whether the culture method and/or SILAC labeling are toxic for the embryonic brain, which could confound further analyses. Cortex proteomics data from embryos cultured with SILAC for 8 hours was compared to freshly dissected cortex from uncultured embryos (**Fig. 1f**). Analysis of 80 gene ontology (GO) pathways encompassing a wide range of toxic responses demonstrated minimal enrichment in cultures at any developmental stage. Taken together, these data benchmark a method for versatile, comprehensive, time-continuous SILAC proteomics in live embryos.

### Time-resolved protein synthesis in cortical development

Embryo culture at stages from E12.5-E16.5 encompasses the chronology of neurogenesis in the cortex ^1^ (**Fig. 2a**). At E12.5, the cortex is predominantly a stem cell tissue, where a pool of neural progenitors has begun generating post-mitotic daughter neurons. Over the next several days of neurogenesis, neurons migrate away from the progenitor pool and sequentially form layers above. Thus, analysis of the cortex from E12.5-E16.5 represents the transition from a progenitor tissue to predominantly differentiating neurons.

**Fig 2.**
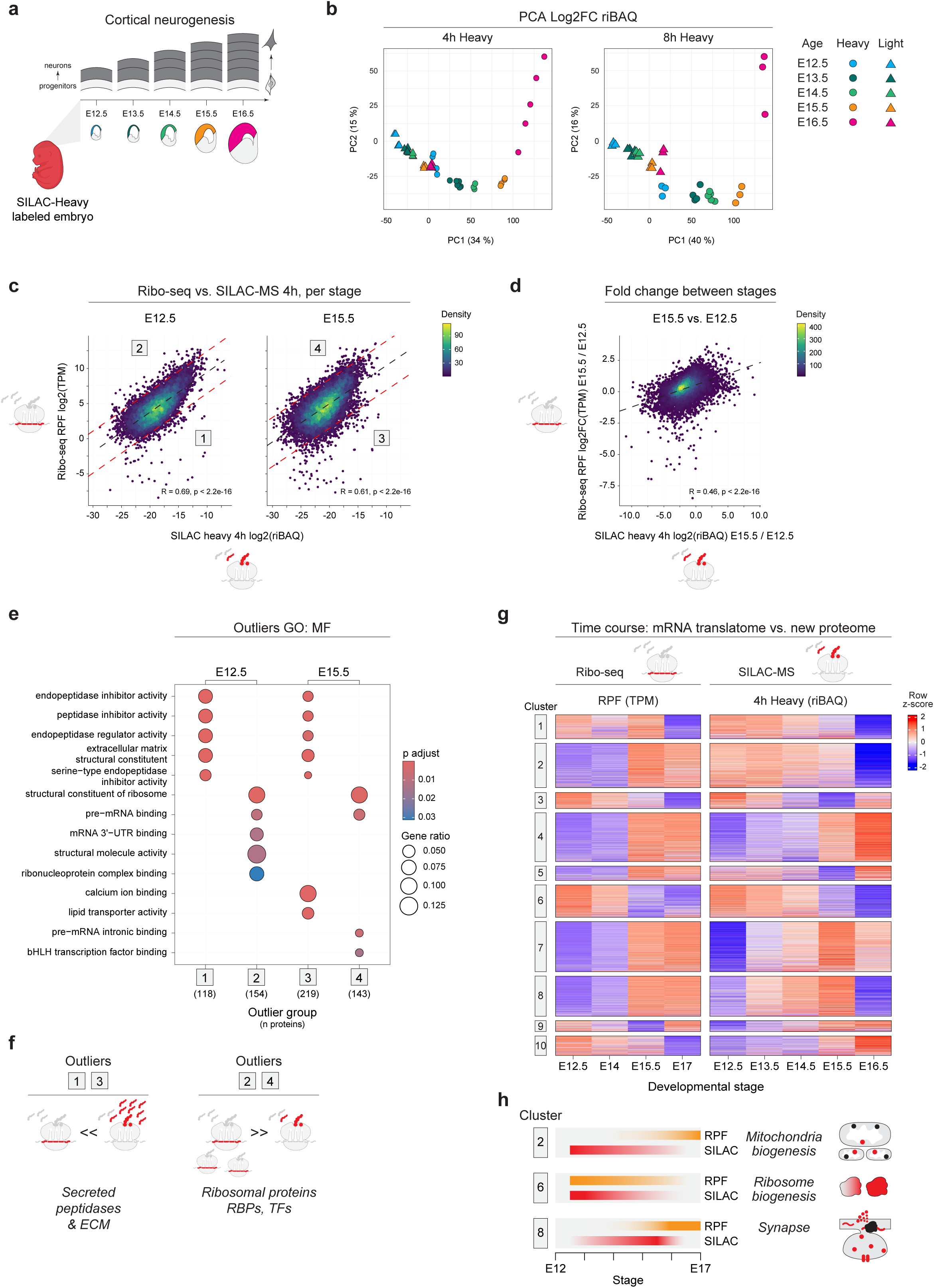
mRNA translation activity and newly synthesized proteins in the developing cortex. **a**, Schematic of cortical tissue analyzed by MEMBRYO and SILAC-MS. From embryonic day E12.5 to E16.5, neural progenitors progressively differentiate into neurons that sequentially form cortical layers. **b**, Principal component analysis (PCA) of heavy and light proteomes from E12.5 to E16.5 cortical tissue after 4 and 8 hours of labeling and SILAC-MS. Variation in the log2 fold change (FC) in relative iBAQ (riBAQ) values between stages is plotted. **c**, Correlation per gene of 4-hour heavy protein labeling and published Ribo-seq data ^16^ from cortex tissue at E12.5 or E15.5. The sum total ribosome protected fragments (RPFs) in the coding sequence were calculated for each gene. The black dashed line is the ordinary least-squares regression; red dashed lines mark ±2 standard deviations from the regression. Numbered boxes (1-4) identify the four outlier groups, cross-referenced in Fig. 2e. **d**, Correlation per gene of the log2 fold change from E12.5 to E15.5 in 4-hour heavy protein and Ribo-seq. **e**, Gene ontology (GO) molecular function (MF) enrichment for each outlier group 1-4 in Fig. 2c. The top enriched terms per group are shown. Dot color indicates the Benjamini-Hochberg adjusted p-value; circle size indicates the fraction of proteins in the group annotated for each term. N proteins per outlier group are shown in parentheses. **f**, Schematic of (e). **g**, Heatmap of row-wise z-score normalized trajectories per gene for Ribo-seq TPM from E12.5-E17 ^16^ (left) and corresponding SILAC heavy protein from E12.5-E16.5 (right). Genes are grouped by k-means clustering (k = 10); cluster labels are shown on the left. **h**, Schematic of (g).

Cortex tissue was analyzed for developmental stage-dependent changes in the newly synthesized heavy proteome vs. old light proteome. Principal component analysis (PCA) was performed on the heavy and light proteomes at each stage from E12.5-E16.5, with either 4 or 8 hours of labeling (**Fig. 2b**). Heavy and light proteomes separated in the first principal component, indicating they yield distinct information (PC1, 34-40%), while the second principal component reflects stage-dependent changes (PC2, 15-16%). Correcting for changes in amino acid availability from E12.5-E16.5 did not significantly impact these findings (**Extended Data Fig. 2a**). Together, these data resolve a sequential time course for the analysis of proteome synthesis and turnover throughout neurodevelopment.

To assess how short (4-hour) SILAC labeling correlates with mRNA translation activity, we next compared the heavy labeled proteome in the cortex to Ribo-seq as an orthogonal measure. Ribo-seq measures ribosome density on mRNA (i.e. “loading” onto mRNA) by sequencing ribosome protected fragments (RPF) as an approximation of translation activity. Mechanisms that regulate translation elongation efficiency ^13^ and acute protein turnover ^14,15^ can lead to distinctions between ribosome loading on mRNA and productive protein output. Thus, comparing RPFs to heavy labeled protein provides insight into how ribosome-mRNA interactions yield varying degrees of protein output.

4-hour SILAC-MS data was compared to developmental stage-matched (E12.5, E15.5) cortex Ribo-seq data published previously ^16^. Heavy protein abundance directly correlates with RPF levels for the corresponding transcripts at E12.5 and E15.5 (R = 0.6 and 0.7, respectively) (**Fig. 2c**). Fold changes in RPF and heavy SILAC between stages E12.5-E15.5 also correlate (R = 0.46) (**Fig. 2d**), albeit less strongly, suggesting generally concordant trajectories of developmental change over time. These data demonstrate that 4-hour SILAC labeling reflects mRNA translation activity differences in the embryonic brain.

Notably, a subset of heavy proteins deviates from trends in the Ribo-seq data, which we annotated as four sets of outliers (**Fig. 2c**). To test whether the outlier proteins are in pathways that might relate to increased or decreased protein output during or subsequent to translation, we performed gene ontology analysis of the outliers (**Fig. 2e-f**). Outlier proteins with high heavy labeling and low RPFs are enriched for secreted peptidase inhibitors and extracellular matrix, which have been characterized as particularly stable, long-lived proteins in brain tissue^17^. In contrast, ribosome-associated proteins are characterized by high RPFs and low heavy protein. Co-translational protein degradation mechanisms have been shown to limit ribosomal protein overproduction, which can compromise cell fitness and survival ^18–21^. Interestingly, some basic helix-loop-helix (bHLH) transcription factors and mRNA-binding proteins demonstrate a similar pattern to ribosomal proteins at E15.5 – high RPFs and low heavy protein – that could likewise reflect rapid protein turnover, and prevent the uncontrolled activation or repression of transcription ^22^. Thus, SILAC proteomics reveals layers of information about newly synthesized proteins in the brain that are distinct from Ribo-seq data.

To compare developmental trends in changing ribosome-mRNA interactions with acute protein output throughout neurogenesis, we clustered proteins by both Ribo-seq and 4-hours heavy SILAC data over the full developmental time course (**Fig. 2g**). The trends in Ribo-seq (E12.5-E17) derived from previously published data ^16^, and heavy SILAC protein (E12.5-E16.5) from this study are in broad agreement for several groups of proteins, while some follow distinct trends. In cluster 2, gene ontology analysis showed that Ribo-seq reads increase while new protein output decreases for mitochondrial and metabolic pathways (**Fig. 2g-h** and **Extended Data Fig. 2b**). Mitochondria undergo fission at later stages as cortical progenitors differentiate into neurons ^23^, when we observe mitochondrial proteins are highly translated but the newly synthesized proteins are depleted. Similarly in cluster 8, newly synthesized synaptic proteins are acutely depleted at E16.5 despite ongoing translation of the corresponding mRNAs (**Fig. 2g-h** and **Extended Data Fig. 2b**). The first cortical synapses are formed at E15.5 by thalamic axons ^24^, which secrete factors that stimulate translation activity ^25^. Shortly after, these prenatal synapses are disassembled and remodeled ^24^, which may be captured in the SILAC data as acute protein depletion. In contrast to clusters 2 and 8, the trends in Ribo-seq and SILAC-MS are concordant for ribosome biogenesis proteins in cluster 6 (**Fig. 2g-h**). Ribosome biogenesis is strongly downregulated at mid-neurogenesis by a decrease in ribosomal mRNA translation efficiency ^16^, which is reflected in ribosomal protein output in our data. Taken together, tracking protein synthesis activity in the brain uncovers key features of post-transcriptional and post-translational regulation where protein output deviates from trends in transcript regulation.

### Changing protein turnover activity diverges from transcription regulation in the cortex

Comparing the new to the old proteome encoded as heavy and light, respectively, reveals both protein synthesis and degradation dynamics ^2–4^. The newly made heavy fraction of the total (i.e. heavy plus light) estimates a protein’s synthesis, while the comparison of heavy to light fractions reflects protein turnover (synthesis and degradation combined). We performed these calculations with the SILAC proteomics data at each stage of cortical development (**Fig. 3a**). The heavy fraction of the total proteome, and the ratio of heavy to light, both show a sequential and progressive decrease in the cortex from E12.5-E16.5. The progressive decrease in global protein synthesis activity may be related to the decrease in ribosome production concurrently ^16^ (**Fig. 2g-h**). These data suggest new protein synthesis, and related protein turnover, globally decline from early to late cortical development.

**Fig 3.**
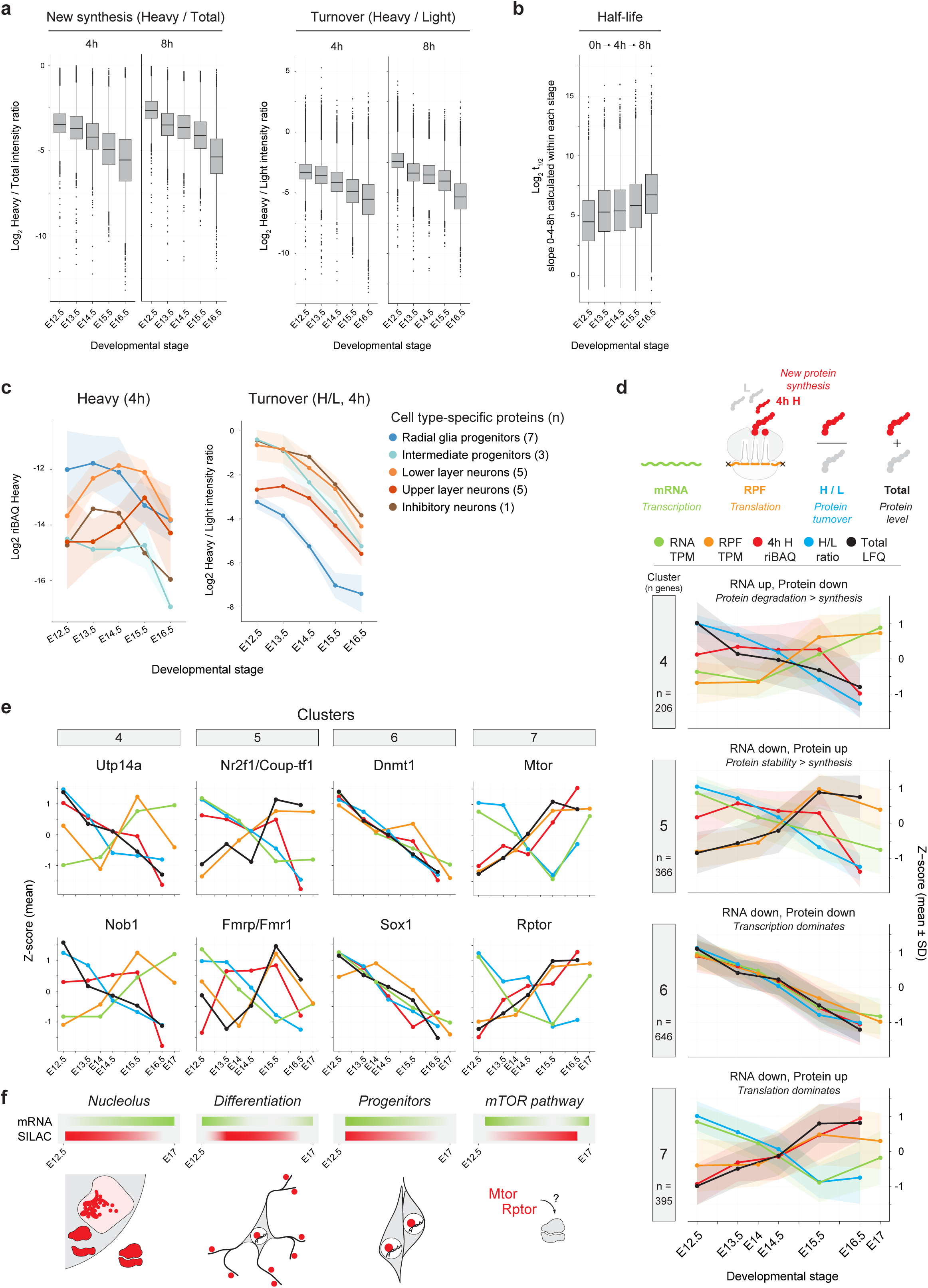
Proteome kinetics and post-transcriptional regulation across cortical neurogenesis. **a**, The distribution of newly synthesized protein (heavy/total), and protein turnover (heavy/light), after 4 and 8 hours of SILAC labeling at each stage. **b**, The distribution of protein half-life (t1/2) calculated from heavy labeling for 0, 4, and 8 hours at each stage. **c**, Trajectories of protein synthesis and turnover activity (4 hours) across developmental stages for marker proteins characteristic of cortical cell types. Marker proteins were defined from published gene sets ^26–28^. Among the proteins characteristic of each cell type are Pax6 for radial glia progenitors, Tbr2/Eomes for intermediate progenitors, Bcl11b/Ctip2 for lower layer neurons, Satb2 for upper layer neurons, and Arx for inhibitory neurons (see **Methods** for full list). **d**, Trajectories of gene expression across developmental stages measured by RNA-seq (green), ribosome protected fragments (RPF) from Ribo-seq (orange), new protein synthesis (heavy SILAC 4-hours, red), protein turnover (heavy/light, blue), and total protein (heavy + light, black). Genes were clustered by similar trajectories between measurements (**Extended Data** Fig. 3b**-c**), and shown are representative cluster numbers 4, 5, 6, and 7 (k = 11 total clusters, N genes per cluster are shown). RNA-seq and Ribo-seq were derived from previously published data ^16^. **e**, Selected genes within the representative clusters, plotted as per (d). **f**, Schematic of (e).

We reasoned that decreasing protein turnover may reflect a shrinking population of progenitors that transition to post-mitotic neurons in the cortex. As cells divide, the pre-existing protein pool is reduced to approximately half. If the proportion of post-mitotic cells increases over time, so would the apparent half-life of the proteome ^2^. To more precisely model protein half-life at each stage, we analyzed the cortex proteome within each developmental stage after 0, 4, and 8 hours of embryo culture, and analyzed its trajectory of change. We focused only on proteins whose total levels are stable within each 8-hour window to minimize non-steady-state effects. Results indicated that proteome half-life progressively increases at each stage (**Fig. 3b**), consistent with an increasing proportion of post-mitotic neurons in the cortex.

To further test how changing cell types contribute to global trends in proteome synthesis and turnover in cortical tissue, we first estimated the relative proportion of cell types in the cortex at each stage from published single-cell RNA-seq data ^26^ (**Extended Data Fig. 3a**). Neural progenitors (radial glia) decline from 57% at E12.5 to 18% at E16.5, whereas their daughter pyramidal neurons increase from 14% to 56%. Then, we analyzed proteins defined by published gene sets ^26–28^ as characteristic of different cortical cell types, and tested whether these specific proteins are differentially regulated by synthesis or turnover in cortical tissue (**Fig. 3c**). The synthesis of proteins associated with radial glia progenitors (eg. Pax6) progressively decreases, while those associated with early lower layer (Bcl11b/Ctip2) and late upper layer (Satb2) neuron differentiation increases, in agreement with their changing proportions in the cortex (**Extended Data Fig. 3a**). In contrast, proteins characteristic of all cell types transition from high to low turnover over time. Thus, some cell type-specific proteins are synthesized selectively, in the context of a global decrease in proteome turnover in the developing cortex.

To trace how multiple steps of gene expression regulation direct ultimate protein output in the cortex, we integrated RNA-seq, Ribo-seq, and SILAC proteomics information per gene over the time course of neurogenesis (**Extended Data Fig. 3b**). Genes were clustered by similar patterns of regulation, which yielded eleven distinguishable clusters (**Extended Data Fig. 3c**), a subset of which are shown in **Fig. 3d**. Genes in cluster 4 are expressed with transcripts (RNA-seq) and their ribosome loading (Ribo-seq) increasing over time, while newly synthesized proteins (4h SILAC), protein turnover, and total protein output decreases. These trends taken together suggest that protein degradation may become greater than synthesis, which is characteristic of nucleolar proteins like Utp14a and Nob1 as shown in **Fig. 3e**. In cluster 5, increasing protein stability may support the accumulation of total protein despite decreasing transcripts and heavy protein output, which is characteristic of genes like Nr2f1 (Coup-tf1) ^29^ that differentiate cortical areas, and Fmrp (Fmr1) ^30,31^ that regulates neuronal differentiation and synaptogenesis. Increasing protein output despite decreasing transcript levels also regulates expression of the mTOR pathway in cluster 7 (Mtor, Rptor ^32^), while neural progenitor genes like Dnmt1 ^33,34^ and Sox1 ^35,36^ in cluster 6 show high concordance between transcript and protein levels. These data highlight that protein synthesis and turnover represent unique cellular states that can diverge substantially from states of transcription.

### Spatial SILAC proteomics analysis of the developing brain

The cortex has a complex architecture with layers of progenitors and differentiating neurons (**Fig. 2a**), and thus resolving proteome synthesis and turnover for different cell types is a challenge in bulk cortical tissue. Spatial proteomics resolves cell type-specific proteomes in complex tissue architectures ^37^, but it has not yet been combined with SILAC labeling *in vivo*. Building upon the SILAC labeling depth of MEMBRYO, we aimed to resolve regional and cell type-specific proteome synthesis and turnover in the embryonic brain.

The E14.5 embryo was labeled with heavy SILAC for 8 hours, and forebrain hemispheres sliced coronally for spatial proteomics (**Fig. 4a**). We first analyzed regions of the forebrain by laser microdissection of the tissue in a grid encompassing the cortex, basal ganglia, and thalamus. The cortex proteome demonstrated the highest SILAC labeling consistent with ongoing neurogenesis at this stage, in contrast to the other regions where neurogenesis is largely complete ^24,38,39^ (**Fig. 4a**). We next focused on the cortex, and progressed to higher spatial resolution in circular areas ranging from 1,000-30,000 μm^2^ in tissue sections 10 μm thick using an ultra-low input spatial proteomics workflow ^40^ (**Fig. 4b**). Tissue volume in this range encompasses approximately 1 to 100 brain cells, depending on the cellular niche ^41^. At a spatial resolution of 5,000-10,000 μm^2^, the progenitor niche in the ventricular zone (VZ) can be distinguished from differentiating post-mitotic neurons in the cortical plate (CP). Proteome depth increases in area-dependent manner, reaching ∼1,000 heavy and 3,500 light at 30,000 μm^2^ following laser microdissection (**Fig. 4c**), providing a platform for spatial SILAC proteomics analysis of cellular niches in the brain.

**Fig 4.**
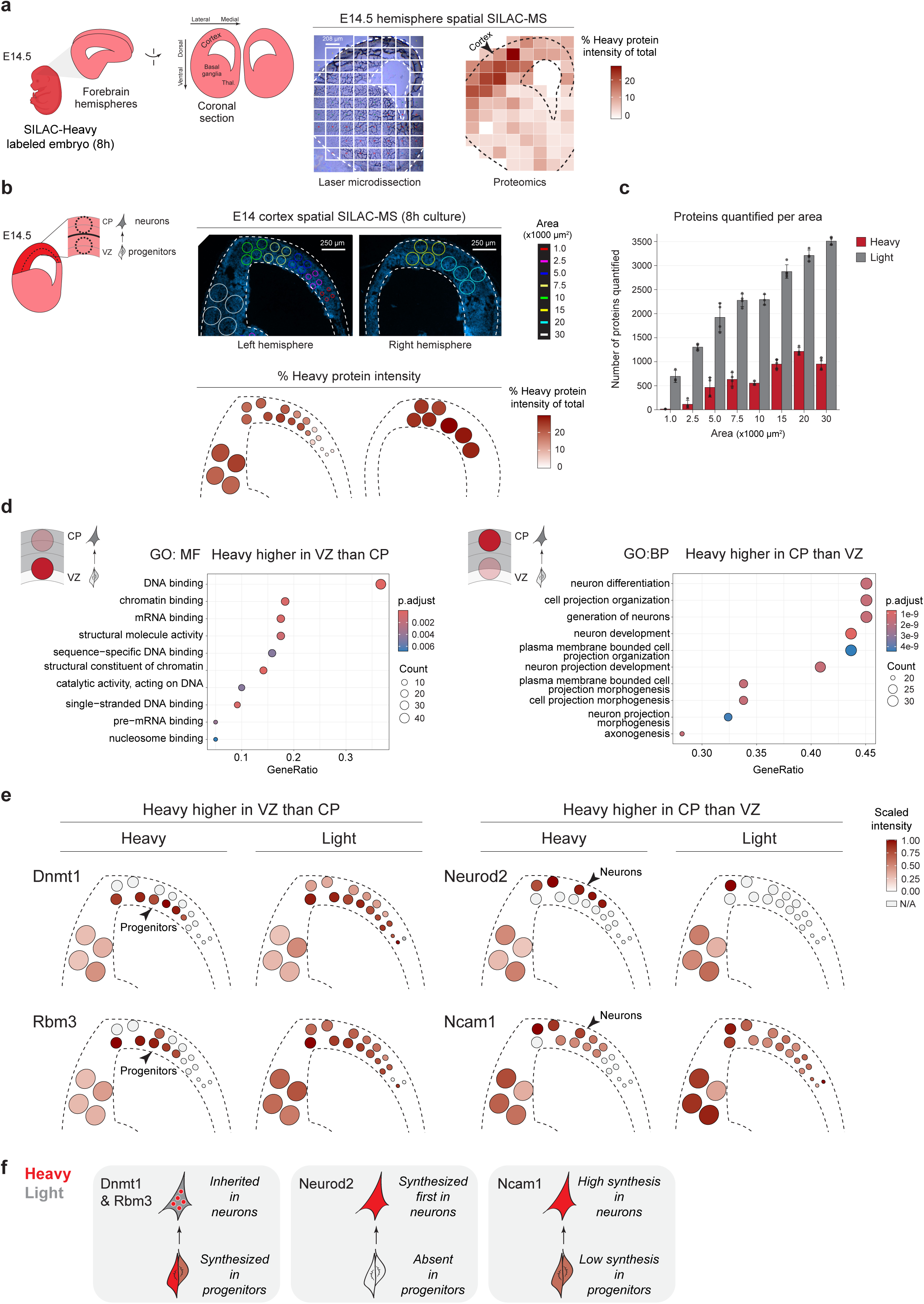
Spatial SILAC proteomics analysis of forebrain regions and cortical layers. **a**, Schematic of an E14.5 embryo heavy labeled for 8 hours, followed by coronal sectioning of the forebrain (left). Spatial map of the laser microdissection grid analyzed by SILAC proteomics, and the percent of total protein intensity that is heavy labeled in each square (right). Newly synthesized proteins are enriched in the dorsal cortex where neurogenesis is ongoing at E14.5, in contrast to ventral regions where neurogenesis is complete at this stage. **b**, Higher spatial resolution laser microdissection of cortical layers in two E14.5 cortical hemispheres, in circles of varying size (top). The percent of total protein intensity that is heavy labeled in each circle (bottom). **c**, Mean number of proteins identified (± SD) per circular area of cortex in the labeled (heavy, red) and unlabeled (light, grey) channels. **d**, Gene ontology (GO) molecular function (MF) enrichment of proteins with higher heavy labeling in ventricular zone (VZ) progenitors plotted on the left, or biological process (BP) enrichment of proteins with higher heavy labeling in cortical plate (CP) neurons on the right. **e**, Spatial map of heavy or light proteins characteristic of progenitors in the VZ (Dnmt1, Rbm3), or differentiating neurons in the CP (Neurod2, Ncam1). The color intensity for each circle is scaled separately per-protein, and for the heavy or light channels. **f**, Schematic of (e).

VZ progenitors and CP neurons were analyzed for distinctions in their heavy proteomes, which represent differentially active molecular pathways (**Fig. 4d**). Consistent with their identities, progenitors synthesize new DNA-binding and chromatin-associated proteins in the VZ, while neurons activate axon projection pathways during differentiation in the CP. Comparison of the heavy and light proteomes in the VZ and CP led to interesting distinctions in synthesis and/or turnover for specific proteins in neurodevelopment and disease pathways (**Fig. 4e**). Dnmt1 and Rbm3 heavy proteins are predominantly synthesized in cortical progenitors in the VZ, and the previously synthesized light proteins are likely inherited in their daughter neurons where synthesis is low (**Fig. 4e-f**). Dnmt1 methylates DNA in neural progenitors ^33,34^, and Rbm3 is a neuroprotection factor during brain injury ^42^, whose cell type-specific mechanisms of action remain unclear in neurodevelopment. In contrast, Neurod2 is a transcription factor that drives differentiation of post-mitotic neurons ^43,44^, which we observed as heavy protein first synthesized in the CP (**Fig. 4e-f**). Ncam1 is a cell surface receptor in both progenitors and neurons with heavy protein likewise elevated in the CP, consistent with its role in cell migration, neurite outgrowth, and synaptogenesis ^45^ (**Fig. 4e-f**). Thus, spatial SILAC proteomics reveals the timing and location of protein dynamics for key neurodevelopmental milestones, which includes tracking proteins that are synthesized in progenitors that are inherited by their daughter neurons.

### Single-cell SILAC proteomics of the developing brain

Single-cell analysis is pushing the detection limits of proteomics, revealing previously hidden diversity in the proteomes of individual neurons ^46,47^ and cell state-dependent protein dynamics^48^. We next leveraged SILAC labeling in the embryo to investigate how protein synthesis and turnover vary across individual cortical cells.

E14.5 embryos were labeled with medium-heavy SILAC (Arg6, Lys4 - “M”) for 8 hours, followed by dissociation and FACS into 384-well plates (**Fig. 5a**). To ensure robust quantification of labeled and unlabeled peptides in cortical cells, the wells were pre-loaded with a heavy (Arg10, Lys8 - “H”) SILAC spike-in proteome derived from N2A cells that were fully labeled in culture. N2A cells are a neuroblastoma cell line with features of both progenitors and differentiated neurons ^49^, and thus are a comprehensive proteome reference. A similar single cell spike-in SILAC (scSiS) strategy was employed previously to improve proteome coverage and quantification accuracy ^48,50^. An additional normalization for global variations in cell size were likewise implemented as previously ^48^. Protein synthesis is estimated by the ratio of medium-heavy to heavy spike-in (M/H), while turnover is estimated by the ratio of medium-heavy to light (M/L). We analyzed 132 cortical cells that demonstrated robust proteome quantification, with around 1000 proteins measured per cell. 496 proteins were quantified in both light and medium-heavy channels per cell on average (**Fig. 5b**).

**Fig 5.**
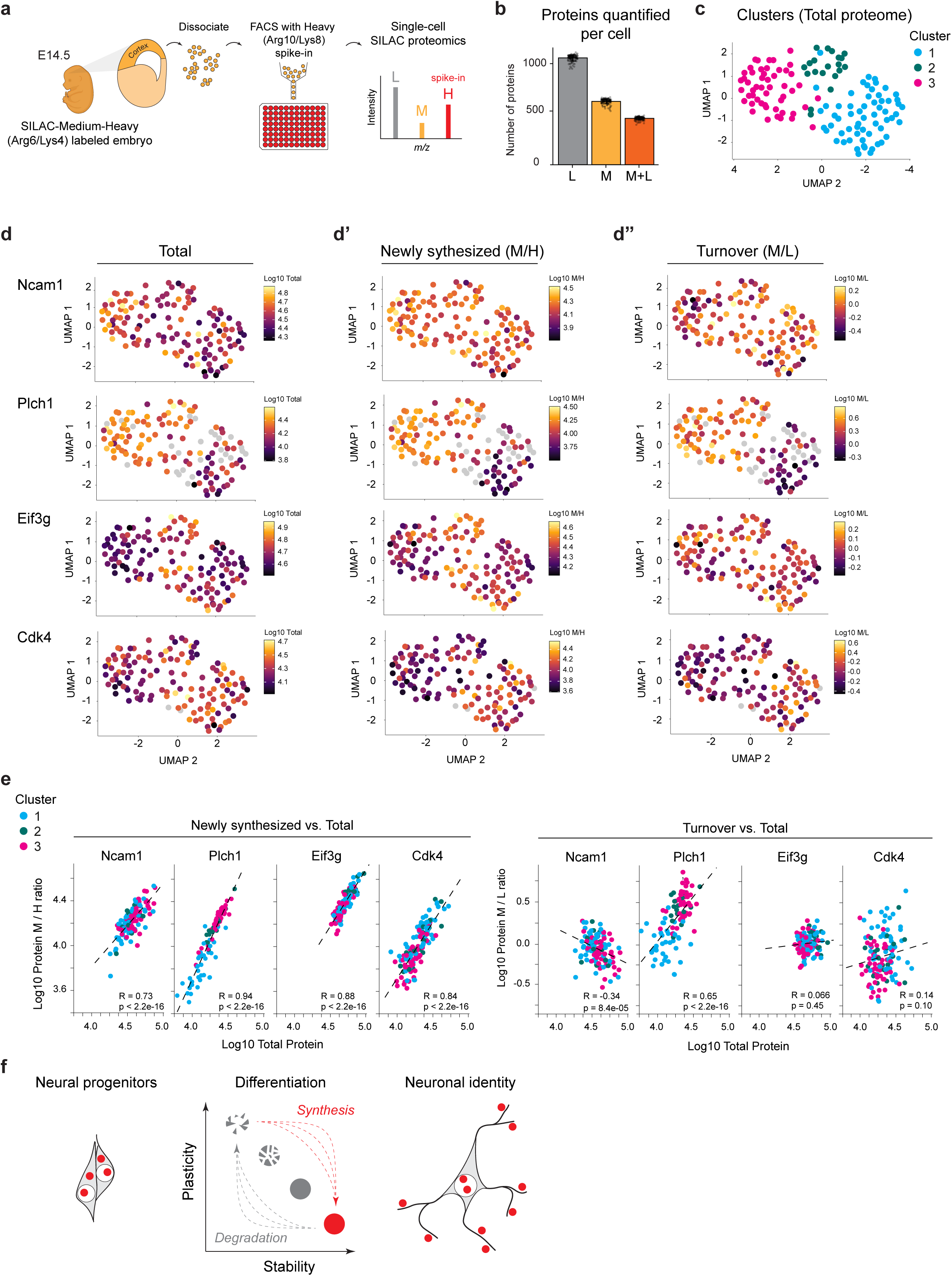
Single-cell SILAC proteomics analysis of protein synthesis and turnover in cortical progenitors and neurons. **a**, Schematic of E14.5 embryos medium-heavy SILAC (M) labeled for 8 hours, followed by the dissociation of single cortical cells, and sorting into multi-well plates. Wells were pre-loaded with a fully heavy-labeled (H) proteome spike-in for normalization and quality control ^48^, and 132 cells were analyzed by single-cell proteomics. **b**, Number of proteins quantified light (L), medium-heavy (M), or in both channels (M+L) per cell. Points represent individual cells (n = 132). **c**, UMAP representation of cells clustered by intensities of their total (M+L) proteome. Cells are colored according to the three distinct clusters identified by shared nearest neighbor (SNN) analysis of the median total intensity per protein per cell. **d**, UMAP from (c) with cells colored by total intensity of selected proteins Ncam1, Plch1, Eif3g, and Cdk4. Color is scaled per-protein. By comparison, cells are shown colored by the scaled intensity of (**d’**) newly synthesized (M/H) protein or (**d”**) protein turnover (M/L). The colors intensity for each cell is scaled separately for each protein and M/H or M/L. Cells without detection of the protein are colored grey. **e**, The correlation of newly synthesized (M/H) protein (left) or protein turnover (right) vs. total protein for Ncam1, Plch1, Eif3g, and Cdk4 per-cell. Each point is a cell colored according to the three distinct clusters annotated as in (c). The black dotted line is the ordinary least-squares regression. Spearman correlation coefficients and associated p-values are shown. **f**, Schematic of how protein synthesis and degradation may direct neuronal differentiation through the selective stability and plasticity of the proteome.

The total proteome of each cell was analyzed in a uniform manifold approximation and projection (UMAP), which yielded 3 distinct clusters of cells by proteome similarity (**Fig. 5c**). Onto this UMAP, we then plotted the total levels of neuronal differentiation and signaling proteins (**Fig. 5d**). Ncam1 and Plch1 are elevated in cluster 3, indicative of a more differentiated neuronal state ^45,51,52^. In contrast, Cdk4 is elevated in cluster 1, which is associated with proliferating cortical progenitors ^53^. Translation initiation factor Eif3g is elevated in cluster 2, which may reflect the translation of mRNAs as neural progenitors transition to differentiation ^54^. Together, these data suggest a sequential transition from proliferative to differentiated neuronal states in clusters 1 to 3, respectively.

We next leveraged the SILAC labeling to investigate how protein synthesis and turnover vary across individual cortical cells. Newly synthesized protein abundance (M/H) was elevated in the same cell clusters enriched for total Ncam1, Plch1, Cdk4, and Eif3g (**Fig. 5d’**). However, the relative abundance of newly synthesized to pre-existing proteins (M/L) exhibited distinct patterns across the four proteins (**Fig. 5d”**). To characterize these relationships, we correlated newly synthesized protein abundance and M/L ratios with total protein abundance across individual cells (**Fig. 5e**). While newly synthesized protein abundance was positively correlated with total abundance for all four proteins, M/L ratios exhibited positive, negative, or no correlation with total abundance, revealing distinct patterns of protein renewal across cellular states. Ncam1 and Plch1 were both enriched in the more differentiated neuronal population (cluster 3), yet exhibited opposing labeling dynamics. For Ncam1, increasing total abundance was accompanied by elevated synthesis but a declining M/L ratio, indicating an increasing relative contribution from the pre-existing protein pool. In contrast, Plch1 exhibited increasing synthesis and M/L ratios with total abundance, indicating a greater contribution from newly synthesized protein. These opposing patterns suggest that Ncam1 accumulation is associated with the preferential retention of pre-existing protein, whereas Plch1 accumulation is associated with increased protein renewal. Eif3g and Cdk4 exhibited relatively constant M/L ratios across cellular states, suggesting that their newly synthesized and pre-existing pools scale proportionally with total abundance. Together, these findings reveal protein-specific differences in synthesis and renewal across individual cortical cells, indicating that neuronal differentiation emerges from a balance of proteome plasticity and stability (**Fig. 5f**).

## Discussion

MEMBRYO advances our understanding of how the molecular landscape of a cell changes as it acquires its identity during development. We find that protein synthesis and degradation dynamics during cortical neurogenesis reveal aspects of cellular state that are not captured by corresponding transcriptional changes, with potential implications for neuronal differentiation and neurodevelopmental disorders. For example, Fmrp (Fmr1), Mtor, and Rptor are key regulators of proteostasis implicated in neurodevelopment and autism spectrum disorders ^55^. We identify developmental stages at which their protein abundance increases despite declining transcript levels, with SILAC measurements implicating regulation at the level of protein synthesis and degradation. These findings extend previous analyses of transcription and translation in the developing cortex ^1,25,26,28,39,56–58^, including recent studies highlighting the distinct, multilayered dynamics of the transcriptome and proteome ^16,47,59^. By extending metabolic labeling to spatial and single-cell proteomics, MEMBRYO further resolves protein dynamics within complex tissues and individual cells. This enables the identification of proteins such as Dnmt1 and Sox1 that are synthesized in neural progenitors and may persist into their neuronal progeny. We also find that the turnover of Ncam1, a neuronal cell adhesion protein synthesized in progenitors ^45^, decreases during differentiation. Features of neuronal identity and connectivity like cell adhesion may be partly consolidated by the regulation of protein stability, in contrast to the phosphoinositide-signaling enzyme Plch1 that exhibits increasing fractional renewal during differentiation. Interestingly, activity-dependent Ncam1 turnover has been implicated in long-term synaptic remodeling postnatally ^60^, raising the possibility that Ncam1 stability also functions in the structural plasticity of mature circuits. Together, these findings suggest that neuronal identity emerges through two interconnected layers of regulation: selective protein synthesis from the transcriptome and selective preservation of the pre-existing proteome.

Beyond these biological insights, MEMBRYO expands the capabilities of *ex utero* embryo culture ^9,10^ by integrating metabolic labeling and extending its application into the third trimester of mouse gestation, encompassing critical periods of brain development. In principle, this approach could be adapted to other models of development and disease, including stem cell-derived embryo models such as blastoids and gastruloids, as well as organoid systems ^61–63^, although modifications to labeling conditions and culture requirements may be necessary. The modularity of embryo roller culture also offers opportunities to introduce additional metabolic probes commonly used in conventional cell culture, potentially enabling measurements of other molecular processes, such as nascent RNA synthesis ^4,5^. Here, we established SILAC labeling as a first application, providing a foundation for extending metabolic tracing beyond proteins to other molecules.

Previous studies employing 4sU and SILAC labeling in cultured cells established approaches for measuring gene expression kinetics, including RNA and protein synthesis and turnover ^4,5,48^. SILAC and click chemistry-based metabolic labeling subsequently enabled investigation of protein dynamics in primary neurons, including local protein synthesis in neuronal processes ^64–66^. *In vivo* SILAC approaches have measured protein turnover in adult mice through the administration of isotope-labeled diets over days to weeks ^3,17,67^. By contrast, MEMBRYO enables continuous metabolic labeling in live embryos over hours, providing access to acute protein synthesis and turnover dynamics during development. Comparing cortical protein turnover measured by MEMBRYO with published measurements from the adult mouse cortex ^17^ revealed only modest correlations between individual proteins (**Extended Data Fig. 4**). These differences may reflect distinct labeling strategies or developmental regulation of protein turnover. MEMBRYO also advances beyond previous approaches involving the injection of isotopically labeled amino acids into embryos ^7^, which achieved relatively low labeling efficiency (∼10% of the proteome) and lacked continuous isotope delivery. Although MEMBRYO enables labeling of nearly the entire detectable proteome in bulk tissue analyses, achieving comparable coverage at spatial and single-cell resolution will require further improvements in mass spectrometry sensitivity. Thus, MEMBRYO advances high-efficiency metabolic labeling beyond cultured cells into the embryo at previously inaccessible developmental stages.

Understanding how molecular states give rise to cellular phenotypes remains a central challenge in developmental and evolutionary biology. By tracing when and where proteins are synthesized and how long they persist, MEMBRYO links molecular history to cellular identity. This may be particularly relevant to evolutionary differences in developmental timing, as protein half-lives have been reported to be longer in human than in mouse neurons ^1,68–70^, potentially contributing to species-specific differences in neuronal maturation. More broadly, the ability to distinguish newly synthesized proteins from those inherited from earlier cellular states raises fundamental questions about how cells integrate their molecular past with their present functions and future developmental trajectories. MEMBRYO opens a window into this temporal dimension of the proteome, providing a framework for investigating how protein synthesis, degradation, and inheritance shape the diverse cellular identities that emerge during embryonic development.

## Methods

### Mice

Wild-type CD1 mice (Mus musculus) were used for all experiments and housed in the animal facilities of the Max Planck Institute for Molecular Genetics. All procedures were conducted in accordance with approval from the Landesamt für Gesundheit und Soziales (LaGeSo) Berlin (protocol G0137/23). Experiments were performed on embryos collected between E12.5 and E16.5; embryonic day 0.5 (E0.5) was defined as the day a vaginal plug was detected. Embryos of both sexes were analyzed without distinction.

### Whole Embryo Roller Culture

Pregnant dams were euthanized at the indicated embryonic stage and embryos were rapidly retrieved from the uterine horns. All dissections were carried out at 37°C in pre-warmed methionine-depleted DMEM (Genaxxon, C4048) on a 37°C heating pad to prevent temperature-induced stress and preserve embryo viability. Embryos were handled exclusively in depleted DMEM to rinse away blood and initiate methionine depletion. They were first kept with the placenta and yolk sac intact, then transferred into fresh depleted DMEM. The yolk sac was opened near the head region, where vascularization is minimal, to reduce the risk of damaging blood vessels or inducing clot formation. Embryos were then released from the yolk sac, the placenta was removed, and embryos were transferred once more to fresh depleted DMEM to further remove residual blood.

For roller culture, embryos were placed into custom-made glass bottles mounted in a BTC Rotating Bottle Culture Unit (Cullum Star LTD, BTC02). Culture media were equilibrated beforehand by gassing with carbogen (5% CO₂/95% O₂; Air Liquide, P3750S10R5A001) for approximately 45 min at 37°C. Embryos were cultured for 4h or 8h at 37°C with continuous rotation and a constant carbogen flow (0.15 LPM; 95% O₂/5% CO₂).

Media formulations were as follows. SILAC media: depleted DMEM (Genaxxon, C4048) to volume, 10% dialyzed FBS (Gibco, A3382001), Sodium Pyruvate 1X (Gibco, 11360-039), GlutaMAX 1X (Gibco, 35050-038), L-Arginine 13C 15N 4 mM (Silantes 201603902) or L-Arginine 13C (Silantes, 201203902) 4 mM, L-Lysine 13C 15N 8 mM (Silantes 211603902) or 4.4’.5.5’-D4-L-Lysine (Silantes, 211103913) 8 mM, L-Methionine 200 µM (Sigma-Aldrich, M5308). Replenished control media: depleted DMEM (Genaxxon, C4048) to volume, 10% dialyzed FBS (Gibco, A3382001), Sodium Pyruvate 1X (Gibco, 11360-039), GlutaMAX 1X (Gibco, 35050-038), L-Arginine 400 µM (Sigma-Aldrich, A5006), L-Lysine 800 µM (Sigma-Aldrich, L5501), L-Methionine 200 µM (Sigma-Aldrich, M5308).

Following incubation, embryos were transferred to 37°C pre-warmed 1x PBS, and examined for visible heartbeat, fluid mobility in the umbilical cord, edema, hemorrhage, or other overt damage. Only embryos with sustained heartbeat and no visible lesions were retained for downstream processing.

### Free amino acid metabolomics

E12.5 and E16.5 embryos were cultured for 4 hours in heavy SILAC medium (4 mM Arg10, 8 mM Lys8). Heart, liver, midbrain, basal ganglia, and neocortex of SILAC-labelled embryos were dissected in PBS and snap-frozen on dry ice. The tissues were then homogenised under denaturing conditions in screw vials (2 mL) with a FastPrep (once for 60 seconds at a speed of 4.5 m x second ^-1^) in a buffer containing 750 µL methanol and 100 µL 0.1% ammonium acetate in water. After a brief centrifugation, the entire lysate was transferred into 15 mL tubes. 2.5 mL methyl-tert-butyl ester (MTBE, Sigma-Aldrich), was added and incubated on a rocking platform, shaking at 1000 rpm at room temperature for 1 hour. This was followed by the addition of 625 µL of LC-MS grade water and again shaking for 10 minutes. The debris was pelleted by centrifugation at a speed of 1,000 g at 4°C for 10 minutes. Both phases were transferred to a new tube and lyophilized. Thereafter, 100 µL of a 1:1 acetonitrile-methanol buffer was added for resolving metabolites, vortexed, and sonicated in a water bath for 10 minutes and centrifuged at 4°C at 3200 rcf for 5 minutes. Supernatants were transferred into new 0.5 mL protein low-binding tubes. A quantity of 45 µL of the supernatant was transferred into the microinjection tubes together with 5 µL of valine D-8 (20 pmol/µL) as internal standard. 10 µL was injected for each LC-MS run in triplicate.

Amino acids were separated on an LC instrument (1290 series UHPLC; Agilent), coupled online to a triple quadrupole hybrid ion trap mass spectrometer QTrap 6500 (Sciex). Data acquisition was performed with an ion spray voltage of 5.5 kV in positive mode, N2 as the collision gas was set to medium, the curtain gas was at 30 psi, the ion source gases 1 and 2 were both at 40 psi, respectively, and the interface heater temperature was set to 350 °C.

Amino acids were separated on a Reprosil-PUR C18-AQ (1.9 µm, 120 Å, 150 x 2 mm ID; Dr. Maisch; Ammerbuch, Germany) at 30°C, on a nonlinear gradient over 39 minutes. The mobile phase consisted of 0.1% formic acid in water (A) and 0.1% formic acid in acetonitrile (B). Multiple reaction monitoring (MRM) was used to measure three transitions per metabolite.

#### Data preprocessing

Peak integration was performed using MultiQuant™ Software v2.1.1 (Sciex). Following extraction, peaks were filtered based on two quality-control criteria. First, transitions were evaluated against validated ion-ratio values: peaks with an absolute deviation >0.15 were excluded, whereas peaks with an absolute deviation between 0.10 and 0.15 were subjected to manual review and curation. Second, transitions were assessed for retention-time consistency; peaks were excluded when the retention-time difference between transitions exceeded 3.

After QC filtering, pseudotransitions were removed. For each amino acid, a single transition was retained by selecting the first transition, except for isoleucine and leucine, for which the second and third transitions were retained, respectively. Peak areas were normalized per sample using a two-step procedure: normalization to the internal standard (Valine D8), followed by normalization to total protein amounts determined by the bicinchoninic acid (BCA) assay.

Poor technical replicates were filtered out based on correlations with replicates, and the median across retained technical replicates was used. The most affected ones were control samples following labelled samples, which is related with layover of previous samples in the liquid chromatography column.

The percentage for the heavy amino acids was calculated from the total pool of amino acids (sum of heavy and light amino acids).

For embryonic ages lacking direct measurements (E13.5, E14.5 and E15.5), the fraction of heavy-labeled amino acid in the intracellular pool was estimated by first-order exponential interpolation on the log scale:

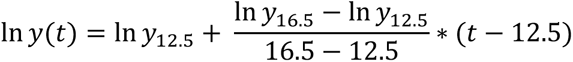

where *y*(*t*) denotes the percentage of heavy amino acid relative to the total pool of the corresponding amino acid at embryonic age *t*.

### Bulk tissue proteomics - SILAC titration

E15.5 embryos were cultured for 4 hours in heavy SILAC medium with the following concentrations: 1x (400 μM Arg10, 800 μM Lys8), 3x, 5x, 10x, 20x or alternatively light control medium. Cortical tissue was microdissected in PBS at 4°C and snap frozen on dry ice.

SILAC-labelled mouse brain tissue samples were processed using a common guanidine hydrochloride-based digestion workflow. Tissue lysates were prepared in 3 M guanidine hydrochloride, 10 mM TCEP, 40 mM 2-chloroacetamide and 100 mM Tris (pH 8.5), heated at 95 °C and sonicated. Protein concentrations were determined by BCA assay, and 30 µg of protein per sample was used for digestion. Lysates were diluted with 10% acetonitrile and 25 mM Tris (pH 8.5) to reduce the guanidine hydrochloride concentration to approximately 1 M and incubated with Lys-C at an enzyme-to-protein ratio of 1:50 for 3.5–4 h at 37 °C with shaking. Samples were subsequently diluted to approximately 0.5 M guanidine hydrochloride and digested overnight with trypsin at an enzyme-to-protein ratio of 1:50 at 37 °C.

Digests were acidified with formic acid, clarified by centrifugation and desalted using Pierce C18 spin columns. Peptides were eluted with 60% acetonitrile and 0.1% formic acid. Approximately 90% of each eluate, corresponding to ∼27 µg was separated into four strong cation-exchange fractions using fractionation buffers 3, 4, 5 and X. SCX fractions were dried by vacuum centrifugation and reconstituted in 5% acetonitrile and 2% formic acid before LC– MS/MS analysis.

#### LC-MS/MS Analysis

3.4 µg per SCX fraction were analysed on a Q Exactive HF mass spectrometer attach to a Dionex UltiMate 3000 Nano LC System following method HF_121_trap_inject_loop_S_v3.

#### Data preprocessing

Raw data was processed with MaxQuant v2.5.0.0 or 2.5.1.0 against mouse protein databases supplemented with common contaminant sequences. Reverse protein sequences were used as decoys, and peptide-spectrum match and protein-level false-discovery rates were controlled at 1%. Peptides of at least seven amino acids were considered. Protein quantification was based on razor peptides, normalized SILAC heavy-to-light ratios and LFQ intensities, and iBAQ calculation was enabled. Oxidation of methionine and protein N-terminal acetylation were included among the modifications used for protein quantification.

Experiments were processed using MaxQuant v2.5.0.0 (Exp 182 and 195) or v2.5.1.0 (Exp 173) with match-between-runs enabled, a matching time window of 0.4 min and an alignment time window of 20 min. A minimum of one peptide per protein was required for experiments 182 and 195, whereas at least two peptides were required for experiment 173. Note: experiment numbers are referenced in the supplied data and code.

MaxQuant proteinGroups.txt outputs were processed in R. Reverse identifications and potential contaminants were removed, protein-group identifiers were reduced to the leading protein identifier, and zero intensity values were converted to missing values. LFQ heavy-channel, LFQ light-channel and normalized heavy-to-light ratio columns were extracted from each experiment for downstream analysis. Proteins were retained when at least two non-missing normalized heavy-to-light ratios were available for the corresponding SILAC analogue and titration concentration.

### Bulk tissue proteomics - SILAC time course

E12.5 to E16.5 embryos were cultured for 4 or 8 hours in heavy SILAC medium (4 mM Arg10, 8 mM Lys8). Cortical tissue was microdissected in PBS at 4°C and snap frozen on dry ice. Sample pellets were resuspended in lysis buffer (1% SDS w/v, 50 mM Tris-HCl pH 8.5) supplemented with cOmplete, Mini, EDTA-free protease inhibitor cocktail (Roche) and immediately boiled at 95°C for 5 minutes. After cooling to room temperature (RT), benzonase (Millipore) was added and samples were incubated for 10 minutes to reduce viscosity. Lysates were then centrifuged at 14,000 rcf for 10 minutes and the supernatant was transferred to fresh tubes. Protein concentration was determined using a BCA protein assay kit (Pierce) and measured at 562 nm on a Multiskan Go plate reader (Thermo Fisher Scientific).

Twenty micrograms of protein per sample was reduced with 10 mM dithiothreitol (DTT; Sigma-Aldrich) at 50°C for 30 minutes, cooled to RT, and alkylated with 22.5 mM iodoacetamide (IAA; Sigma-Aldrich) for 30 minutes in the dark at RT. The alkylation reaction was quenched by addition of 10 mM DTT.

SP3 cleanup was performed using magnetic SpeedBeads (Cytiva; cat. no. 45152105050250 and 65152105050250). Beads were added at a 20:1 bead-to-protein ratio and 50% acetonitrile (ACN) was added to induce protein binding. After 10 minutes of shaking, the supernatant was removed on a magnetic rack and the beads were washed three times with 80% ethanol. Proteins were digested on-bead by resuspending in 50 mM ammonium bicarbonate (ABC) with trypsin (Promega) at a 1:50 enzyme-to-protein ratio and incubating overnight at 37°C. Peptides were recovered from the beads on a magnetic rack, dried by vacuum centrifugation, and resuspended in buffer A at the desired concentration for LC-MS analysis.

#### LC-MS/MS Analysis

Samples were analyzed by data-independent acquisition (DIA) on an Orbitrap Astral mass spectrometer coupled to a Vanquish Neo UHPLC system (Thermo Fisher Scientific). Reverse-phase separation was performed using in-house packed columns (20 cm × 75 μm inner diameter, 1.9 μm ReproSil-Pur C18-AQ silica beads) heated to 50°C and connected to an electrospray ionization source (Thermo Fisher Scientific). Peptides were separated at a constant flow rate of 250 nL/min over a 38-minute gradient. Mobile phase A consisted of 3% acetonitrile and 0.1% formic acid in water, and mobile phase B consisted of 90% acetonitrile and 0.1% formic acid in water (all LC-MS grade; CHEMSOLUTE, Fluka). The gradient was as follows: 2–7% B over 1 minute, 7–20% B over 19 minutes, 20–30% B over 9 minutes, 30–60% B over 3 minutes, followed by a column wash at 90% B for 6 minutes.

Full MS scans were acquired in the Orbitrap at a resolution of 240,000, with a scan range of 380–1100 m/z, RF lens at 55%, normalized AGC target of 500%, and a maximum injection time of 3 ms. DIA scans were acquired with a precursor mass range of 380–980 m/z and an isolation window of 4 Th. HCD fragmentation was performed at a normalized collision energy of 25%. Fragment ion scans covered a range of 150–2000 m/z with an RF lens of 40%, normalized AGC target of 500%, and a maximum injection time of 3 ms.

#### Data Preprocessing

Raw files were analysed using DIA-NN ^71^ v1.8.1 with a spectral library generated from the UniProt database (UP000000589_10090.fasta). The following settings were used: match-between-runs (MBR) off, requantification on, precursor m/z range 400–1000, mass accuracy 10, MS1 accuracy 15, scan window 3 and library generation, ID, RT and IM profiling. The following additional options were applied: --fixed-mod SILAC,0.0,KR,label; --lib-fixed-mod SILAC; --channels SILAC,L,KR,0:0; SILAC,H,KR,8.014199:10.008269; --peak-translation; -- original-mods; --relaxed-prot-inf.

The report.tsv output file output was subsequently processed using the software, stackedLFQ (https://github.com/rkerrid/StackedLFQ, adapted from a similar implementation in ^72^). Briefly, the DIA-NN report.tsv was filtered using the following criteria: Global.PG.Q.Value < 0.01, Precursor.Charge > 1, and Channel.Q.Value < 0.03. Precursors labelled as contaminants were removed.

Precursor.Quantity values of the remaining SILAC pairs were summed to calculate total ion intensity and used to derive precursor heavy-to-light (H/L) ratios. Protein group level ratios were calculated as the median of precursor SILAC ratios, requiring a minimum of two ratios per protein group per run. Proteins were filtered by a minimum of 2 replicates with ratios per embryonic age and incubation time.

LFQ values was an output of the stackedLFQ package were it was calculated using the directLFQ package ^73^ on the total ion intensity.

The output file ratios.csv was used to calculate the heavy-to-total fraction as follows:

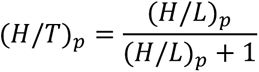

where the subscript *p* denotes protein-specific values.

The output file ratios.csv was used to calculate the light-to-total fraction as follows:

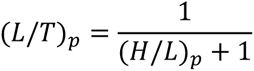

where the subscript *p* denotes protein-specific values.

Samples identified as outliers during quality control were removed before downstream analysis. Proteins were retained when an H/T value was available in at least two biological replicates for the corresponding embryonic age and incubation time.

To account for developmental differences in precursor-pool labelling, precursor heavy quantities were divided by the age-specific intracellular heavy-amino-acid fraction. Heavy-to-light free amino acid normalized precursor ratios were calculated as follows:

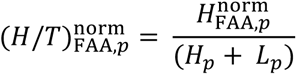

where *H*^norm^ denotes the normalized heavy free–amino-acid measurement used for precursor *p*, and *H_p_* and *L_p_*are the corresponding heavy and light signal for precursor *p* from the output file precursors.csv. Proteins ratios were rolled up as the median of the precursors’ normalized ratios.

Intensity-based absolute quantification (iBAQ) values for both heavy and light channels were calculated from precursor-level intensities. Theoretical tryptic peptides were generated *in silico* according to the Keil rule (cleavage C-terminal to Lys and Arg, except when followed by Pro). Only peptides with lengths between 6 and 30 amino acids were considered observable. The iBAQ value for each protein was defined as the sum of its precursor intensities (from precursor.csv file) divided by the number of theoretical observable peptides.

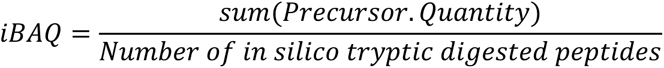

To enable inter-sample comparison, relative iBAQ (riBAQ) values were determined by normalizing the iBAQ of each protein (*p*) to the total iBAQ sum within each sample (*s*):

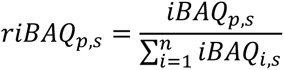

where *n* represents the total number of proteins detected in the sample.

#### Quality control and statistical summaries

Protein counts were calculated as the number of non-missing protein measurements per sample. Unless stated otherwise, bar plots show the mean and standard deviation across biological replicates. Comparisons of 4- and 8-h incorporation distributions were performed using two-sided Wilcoxon rank-sum tests within each embryonic age and analogue, followed by Benjamini–Hochberg correction. PCA was performed on log2-transformed riBAQ values using proteins quantified without missing values in the samples included in each analysis. Variables were centered and scaled before PCA.

#### Estimation of arginine to proline conversion

To test for arginine to proline conversion, we analyzed DDA measurements using MaxQuant (v2.6.8.0) with heavy proline as a variable modification. Peptides were filtered for cases where a heavy SILAC peptide would be quantified in both a light and heavy proline version. For those peptides, the fraction of heavy proline was calculated as 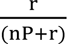 where r is the ratio of intensity of the converted (heavy Proline) form to that of the unconverted form and nP is the number of prolines in the peptide. The data were plotted combined for each stage from E12.5-E15.5; E16.5 did not yield sufficient peptide coverage for estimation.

#### Estimation of peptide labeling efficiency

Labelling efficiency was determined by analyzing DDA measurements with MaxQuant (v2.6.8.0) changing default settings to 3 missed cleavages and avoiding the SILAC settings and rather adding the heavy SILAC amino acids as variable modifications. This allows to check for mis-cleaved peptides with both versions, fully heavy labelled peptides as well as peptides with a mixed combination of light and heavy amino acids. To calculate the labelling efficiency, this formula was used 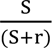 where S is the number of possible SILAC sites (number of arginine and lysine in the peptide summed) and r is the ratio of intensity 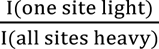. The data were plotted combined for each stage from E12.5-E15.5; E16.5 did not yield sufficient peptide coverage for estimation.

#### GSEA of toxicity pathways

The LFQ output from the software, stackedLFQ (https://github.com/rkerrid/StackedLFQ, adapted from a similar implementation ^72^) after log2 transformation was used to rank proteins in control (uncultured cortex) and SILAC samples. Rank differences were computed as rank_control - rank_silac and were used as input in a Gene set enrichment analysis (GSEA) using the package fgsea, in preranked mode using default settings.

The 80 gene sets used were downloaded from the Molecular Signatures Database (MSigDB) and were related to cell death (necrosis, autophagy and apoptosis), cellular stress (DNA damage, cytotoxicity, and chemical, heat and oxidative stress) and inflammation.

#### Dimensionality Reduction and Procrustes Analysis

To evaluate the impact of the availability of heavy amino acids on the global proteomic profile, we performed Principal Component Analysis (PCA) followed by Procrustes transformation. To ensure a balanced comparison between the standard (*H*/*T*) and the free-amino-acid-normalized 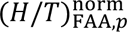 datasets, we restricted the analysis to a core set of proteins consistently quantified across all samples. PCA was performed to extract the principal configurations of the samples in reduced dimensional space. To quantify the concordance between the two normalization approaches, we applied a Procrustes transformation using the vegan R package (https://rdocumentation.org/packages/vegan/versions/1.0-1), which rotates, scales, and reflects the 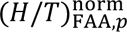 configuration to optimally superimpose it onto the (*H*/*T*) target configuration while minimizing the sum of squared residuals (*M*^2^)

The statistical significance of the Procrustes correlation (*t*_0_) was assessed using a Procrustean randomization test (protest) with 999 permutations. The resulting p-value and sum of squares were used to determine the degree of structural deformation introduced by free-amino-acid normalization. For visualization, the Procrustes-aligned coordinates were plotted, with vectors (arrows) connecting identical samples between the two configurations. This displacement represents the sample-specific shift in multivariate space attributable to the normalization process, categorized by embryonic age and incubation time.

#### Integration of datasets

We utilized our previously published Ribo-seq data from the embryonic neocortex ^16^ for comparison to the SILAC-MS. UniProt, Ensembl and gene-symbol identifiers were mapped using org.Mm.eg.db. TPM of ribosomal protected fragments (RPF) and riBAQ values were merged by common gene (Ensembl ID) and embryonic age. The same values for Riboseq were used to compare them with riBAQ at 4h and 8h. Translation efficiency was calculated as TPM RPF/TPM RNAseq. A linear regression was fit between log2 TPM and log2 riBAQ. Model fit was summarized using the coefficient of determination (R²) and the standard deviation was calculated.

To quantify translational changes between embryonic ages E15.5 and E12.5, log2 fold-changes were computed per gene from the normalized intensity values (TPM/riBAQ). For each gene and timepoint, replicate values of were averaged (median) to obtain a single value per gene per age. Entries with missing values were removed prior to fold-change calculation.

Log2 fold-change was then computed as the log2 ratio of E15.5 and E12.5 values:

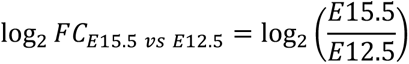

Pairwise Spearman rank correlations were computed using the R function “cor” with pairwise deletion of missing values, such that each correlation coefficient was estimated using all available observations for the corresponding variable pair.

To enable comparison of expression trajectories across experiments, replicate values were summarized by the median (log2 TPM and log2 riBAQ) for each protein, assay and embryonic age and normalized by mean-centering (z-score) separately within each protein and assay across embryonic ages. Only genes with complete data were used for the k-means clustering. K-means clustering was performed with 50 random initializations and a fixed random seed. Ten clusters were used for the RPF–riBAQ-H analysis and 11 clusters for the integrated RNA-seq, RPF, total LFQ, H/L and riBAQ-H analysis, based on inspection of within-cluster sums of squares.

The comparison between the SILAC-MS timecourse and published data was done using processed data provided by E.F.F, derived from published raw data ^17^. Median H/L ratios per protein were used from the stackedLFQ package output (see “Bulk proteomics: DIA Liquid Chromatography with tandem mass spectrometry (LC-MS/MS)”) for our time course and from the MaxQuant output as reported ^17^. Pairwise Spearman rank correlations were computed using the R function “cor” with pairwise deletion of missing values, such that each correlation coefficient was estimated using all available observations for the corresponding variable pair. A linear regression was fit between log2 ratios in our time course and at 5, 14 and 21 days in the Fornasiero et al. 2018 dataset ^17^. Model fit was summarized using the coefficient of determination (R²).

#### Gene Ontology (GO) Analysis

Protein identifiers were mapped to Entrez IDs using the Mus musculus genome-wide annotation database (org.Mm.eg.db, version 3.19.1). To identify overrepresented biological themes, functional enrichment analysis was performed across the three Gene Ontology (GO) domains (Molecular Function (MF), Cellular Component (CC), and Biological Process (BP)) using the compareCluster function in the clusterProfiler R package ^74^. All proteins quantified in this study were used as the background proteome. P-values were adjusted for multiple testing using the Benjamini–Hochberg (BH) procedure. Significant terms were defined by an adjusted p-value < 0.05 and a q-value < 0.2. To reduce semantic redundancy for visualization, enriched GO terms were collapsed using the simplify function with a similarity threshold of 0.7.

#### Kinetics

Protein turnover rates were determined by measuring the incorporation of heavy-labeled amino acids over a pulse-labeling period of 0, 4, and 8 hours. Following the methodology described by Fornasiero et al. 2018 ^17^, the degradation rate (δ) for each protein was estimated by fitting the ratio of heavy-to-total protein (r) to a first-order kinetic model across the three time points. Under steady-state conditions, where protein synthesis and degradation rates are balanced, the fraction of heavy-labeled protein at time t is expressed as:

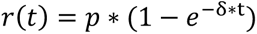

Where *r(t)* is the ratio of heavy-to-total protein at time t, δ is the degradation rate constant and *p* represents the precursor pool enrichment, defined as the fractional abundance of heavy amino acids available for protein synthesis.

#### Estimation of protein half-life

We opted for a strategy that estimates protein half-life within each stage, since protein degradation is difficult to disentangle from a global decrease in protein synthesis that occurs from early-to-late developmental stages. To do so, we calculated the slope of Heavy/Total and Light/Total at 0, 4, and 8 hours of culture at each of the five stages. The analysis was employed only for proteins approximating a steady-state during 8 hours of culture, which we estimated by a linear model and empirical Bayes moderated t-tests fit between the 4h and 8h time points for each developmental stage, using the total abundance per protein. Statistical analysis was conducted using the *limma* R package ^75^. Proteins were defined as steady-state if the adjusted p-value > 0.05 and *log*_2_*FC* < 0.6.

We further attempted to correct for changes in labeling efficiency. The precursor pool enrichment constant used in the model at E12.5 and E16.5 was defined as the heavy-label percentages, calculated from the mean normalized peak areas of free arginine and lysine measured by MS in neocortex, and for E13.5, E14.5 and E15.5 as the interpolated values (see data analysis *Free amino acid Mass Spectrometry*). By incorporating these age-specific *p* values into the kinetic equation, we corrected for the dilution of the SILAC label by endogenous, unlabelled amino acids, ensuring that the calculated degradation rates reflect true protein turnover rather than label availability.

Proteins half-lives (t1/2) were derived from the estimated degradation rates using the formula:

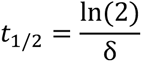

Cortical cell type composition analysis from published single-cell RNA-seq data (Di Bella et al. 2021 ^26^)

Published 10x Genomics single-cell RNA-seq data from mouse cortex were derived from Di Bella et al. 2021 ^26^. Count matrices from E12.5–E16.5 mouse neocortex were imported into Seurat. Genes detected in fewer than three cells and cells with fewer than 200 detected genes were removed during object creation. Cells with at least 6,000 detected genes or at least 20% mitochondrial reads were excluded.

Datasets were merged, library-size normalized and analysed using 3,000 variable genes. Data was scaled and subjected to PCA. Developmental-stage effects were corrected with Harmony using embryonic age as the grouping variable. A shared-nearest-neighbour graph was constructed from the first 30 Harmony dimensions, followed by graph clustering at resolution 0.5 and UMAP visualization.

Cells were annotated using module scores calculated from curated marker sets for radial glia (*Pax6*, *Sox2*, *Vim*, *Nes*, *Slc1a3*, *Fabp7*, *Aldoc, Hes1*, *Hes5*), intermediate progenitors (*Eomes*, *Insm1*, *Sstr2*, *Gadd45g, Btg2, Neurog2*), early (*Bcl11b*, *Fezf2*, *Neurod6*, *Sox5*, *Tbr1*) and late (*Satb2*, *Cux1*, *Cux2*, *Pou3f2*, *Pou3f3, Rorb*) excitatory neurons, inhibitory neurons (*Arx, Gad1, Gad2, Dlx1, Dlx2, Lhx6*), Cajal–Retzius cells (*Reln*, *Lhx5*, *Trp73*, *Calb2*), endothelial cells (*Pecam1*, *Cldn5*, *Esam*, *Kdr*), and microglia (*Cx3cr1*, *Aif1*, *Tmem119*, *P2ry12*, *Csf1r*). Each cell was assigned to the cell type with the highest module score; cells for which all scores were negative were labelled unknown. Cell-type proportions were calculated within each embryonic age after annotation.

The same genes were utilized for plotting the synthesis/turnover of cell type-specific proteins in the SILAC-MS data in **Fig. 3c**; however, Hes1, Hes5, Gadd45g, Btg2, Neurog2, Rorb, Gad1, Gad2, Dlx1, Dlx2, and Lhx6 were not included due to insufficient detection of heavy-labeled peptides for these proteins.

### Spatial SILAC proteomics

#### Mouse E14.5 forebrain hemisphere (square grid at lower spatial resolution)

E14.5 embryos were cultured for 8 hours in heavy SILAC medium (4 mM Arg10, 8 mM Lys8). Brains were microdissected in PBS at 4°C, embedded in optimal cutting temperature (OCT) compound, and frozen on liquid nitrogen. Coronal sections of the forebrain were cryo-sectioned at 100 µm thickness, mounted on DIRECTORTM slides (AMR), and stored at -80°C until processing. Square regions of interest (ROIs; an area of 43,044 µm² per sample) were cut by laser microdissection with the LMD7 (Leica) with a 6.3× objective in brightfield mode at the following settings: power 50, aperture 17, speed 6, specimen balance 3, head current 100%, pulse frequency 800, offset 60. The “Draw + Scan” mode was used. Tissue samples were collected into an individual well of a 96 well plate (twin.tec PCR plate 96 well, semi-skirted, Eppendorf) containing 25 µl of PTS buffer: 12 mM Sodium Deoxycholate (FUJIFILM Wako, 194-08311), 12 mM Sodium N-Lauroylsarcosinate (FUJIFILM Wako, 192-10382) in Tris-HCl pH8.

After LMD, the plate was heated at 95°C for 5 minutes and centrifuged briefly at 500g. For sample digestion, 100 µl of 50 mM Ammonium Bicarbonate (FUJIFILM Wako, 017-02875) and 20 ng of Trypsin (Promega, V5113), were added to each sample. The samples were incubated overnight at 37°C while shaking at 500 rpm. After digestion, 100 µl of ethyl acetate (FUJIFILM Wako, 051-00351) and 20µl of 10% Trifluoroacetic acid (FUJIFILM Wako, 208-02741) were added to the sample. Desalting was performed with SDB-RPS disks (CDS Analytical, 5065-64044) and eluted in 30 µl 5% ammonia (FUJIFILM Wako, 010-03166) 80% acetonitrile (FUJIFILM Wako, 014-00381). After speedvac, samples were resuspended in 0.1% Trifluoroacetic Acid + 0.02% LMNG (Nacalai Tesque, NG310), sonicated and loaded into Evotips as per manufacturer’s instructions.

The Evotip Pure sample was analyzed using an Evosep One system (EVOSEP) equipped with an Orbitrap Exploris 480 mass spectrometer. Evosep One was acquired using the Whisper 20 SPD method (gradient running time, 58 min; flow rate, 100 nl/min). The digested peptides were separated using an IonOpticks Aurora Elite (EV1112) column (5 cm × 75 μm i.d., C18 particle size 1.9 μm) at 35 °C. Mobile phases A and B consisted of 0.1% formic acid and 0.1% formic acid in 100% acetonitrile, respectively. The peptides eluted from the column were analyzed using an Orbitrap Exploris 480 instrument with data-independent acquisition (DIA). The following settings were maintained throughout data acquisition: positive mode; electrospray voltage, 1.8 kV; ion transfer tube temperature, 275 °C; default charge state, 3; orbitrap resolution, 60,000; RF lens, 40%; centroid data. Full MS scans were collected in the range of 450 to 1000 m/z with an AGC target of 300%. MS2 spectra were collected at 200 to 1800 m/z to set an AGC target of 3000% and normalized HCD collision energies of 22, 26, and 30%. The isolation width for MS2 was set to 14 m/z.

Raw files were analysed using DIA-NN v1.8.1 with a spectral library generated from the UniProt database (UP000000589_10090.fasta). The following settings were used: match-between-runs (MBR) on, requantification off, precursor m/z range 200–1800, mass accuracy 10, MS1 accuracy 15, scan window 3 and library generation, ID, RT and IM profiling. The following additional options were applied: --fixed-mod SILAC,0.0,KR,label; --lib-fixed-mod SILAC; --channels SILAC,L,KR,0:0; SILAC,H,KR,8.014199:10.008269; --peak-translation; -- original-mods; --relaxed-prot-inf.

The report.tsv output file output was subsequently processed eliminating the contaminants and proteins with less that 2 peptides and calculating the protein intensities through the median of the precursor.translated intensities of the peptides.

#### Mouse E14.5 cortex (circular regions in cortical layers at higher spatial resolution)

E14.5 embryos were cultured for 8 hours in heavy SILAC media (4 mM Arg10, 8 mM Lys8). Brains were dissected in PBS at 4°C, and fixed overnight in 4% PFA at 4°C while shaking. Fixed brains were embedded in OCT and frozen in liquid nitrogen. Coronal sections were cut using a cryostat at 10 μm thickness and mounted onto PPS Metal FrameSlides (cat. no. 11600294). Sections were thawed in the slide and dried at 37°C for 1h. Slides were immersed in ice-cold 50 mM ammonium formate (cat. no. 09739-500G) to remove residual OCT, then air-dried for an additional 1 h in a fume hood. Sections were stained with DAPI in PBS 1X for 2h at room temperature.

Slides were imaged using the Zeiss Axioscan 7 slide scanner using a 10X objective and 2x2 binning. Regions of interest represented basal ganglia and cortex were delineated anatomically and included technical quadruplicates of areas of 1k (k = 1,000 µm^2^), 2.5k, 5k, 7.5k, 10k, 15k, 20k and 30k in each region. These were manually labeled using the open-source software for digital pathology image analysis QuPath (v0.5.1). Regions of interest were collected by laser microdissection (LMD) on a Leica LMD7 microscope using 20x and 40x objectives operated in brightfield mode, with each replicate into an individual well of a LoBind 384-well plate.

Collected cells were processed for bottom-up LC-MS based proteomics. Acetonitrile (2x10uL) was pipetted to each well to drag samples to the bottom of their wells and evaporated by vaccum-drying centrifugation. For cell lysis, 2 µl of lysis buffer (0.1% DDM, 5 mM TECP, 20 mM CAA, 100 mM TEAB pH 8.5 in LCMS-grade water) was added to each well, shortly centrifuged (2,000 RCF, 1 min) and plate heated at 95°C for 60 min in a thermal cycler (Bio-Rad, 384-well reaction module) at a constant lid temperature of 110°C. Samples were cooled to room temperature and 1 µl of 4ng/μl lys-C added, prediluted in 60% LCMS-grade water 30% ACN and 10% TEAB and incubated at 37°C for 4 hours in the thermal cycler. Subsequently, 1 µl of 6ng/μl trypsin (Promega Trypsin Gold) was added pre-diluted in 80% LCMS-grade water 10% ACN and 10% TEAB and incubated overnight at 37°C in the thermal cycler. The next day, digestion was stopped by vacuum-drying the samples (40min at 60°C). Samples were stored at -20°C until LC-MS instrument availability. Samples were loaded into Evotips as per manufacturer’s instructions. Briefly, Evotips were equilibrated by running through 20 μL Buffer B (0.1% formic acid in 100% ACN) followed by 20 μL Buffer A (0.1% formic acid in LC-MS grade water), and soaking in isopropanol for 10 seconds. Samples were reconstituted in 10 μL Buffer A and transferred into Evotip, flowed through with 20 μL Buffer A and tips filled with Buffer A. Samples were eluted using the Evosep LC system at 40 samples per day (SPD) with WhisperZoom setting into a trapped ion mobility spectrometry quadrupole time-of-flight mass spectrometer (timsTOF Ultra2, Bruker Daltonik GmbH, Germany) with a nano-electrospray ion source (Captive spray, Bruker Daltonik GmbH). Peptides were loaded on a 15 cm IonOptics Aurora Elite HPLC-column (75 µm inner diameter packed with 1.9 µm C18 beads). Column temperature was controlled by a column oven and kept constant at 50°C. Mass spectrometric acquisition was performed in data-independent (diaPASEF ^76^) mode using the default method for long gradients. Ion accumulation and ramp time in the dual TIMS analyser was set to 100 ms each and we analysed the ion mobility range from 1/K0 = 1.6 Vs cm-2 to 0.6 Vs cm-2. The total m/z range was set to 100-1,700 m/z. The collision energy was lowered linearly as a function of increasing mobility starting from 59 eV at 1/K0 = 1.6 VS cm-2 to 20 eV at 1/K0 = 0.6 Vs cm-2. (timsControl software, Bruker Daltonik GmbH).

The data was analyzed as per Bulk tissue proteomics: *Data Preprocessing*.

Protein-group ratio, total, light and heavy LFQ intensity tables were converted to long format and matched to anatomical-region metadata and spatial coordinates. Samples were labelled as outliers and eliminated based on the intensity values and number of proteins following interquartile range (intensity/number of proteins < (Q1 - 1.5 * IQR) or intensity/number of proteins > (Q3 + 1.5 * IQR)).

Differential heavy-channel abundance between ventricular-zone and cortical-plate regions was analysed with proDA. Zero intensities were treated as missing and non-zero intensities were log2 transformed. A proDA model containing anatomical-region and analogue combinations was fitted, and the contrast (VZ_heavy-CP_heavy) was tested. Proteins were classified as differentially abundant when the Benjamini–Hochberg-adjusted (P<0.05) and the absolute log2 difference exceeded 0.6. GO analysis used all proteins included in the proDA result as the experiment-specific background.

### Single cell SILAC proteomics

#### Spike-in preparation

Sample spike-in processing was performed as described in the Methods section of Welter and Mutschler et al. 2026 ^48^, specifically in the subsections “Cell sorting”, “Single-cell sample preparation” and “LC-MS/MS analysis – single-cell measurements”, with minor alterations. The neuroblastoma cell line Neuro2a (N2a) proteome were used as a spike-in. These cells were grown in the same conditions as described in the methods section for 6-8 passages and complete heavy labeling was confirmed by mass spec. Lysis buffer and 1 ng of the digested N2a spike-in were pre-distributed in a 384-well plate, before sorting a single cell suspension of dissociated E14.5 cortex cells into the wells as described below.

#### Single-cell labeling and dissociation from the cortex

Five E14.5 embryos were cultured for 8 hours in medium-heavy (Arg6, Lys4) media in the same conditions and concentration for the heavy labeling experiments above (4 mM Arg6, 8 mM Lys4). Immediately after, single cells were dissociated from microdissected cortex tissue and pooled between the five embryos. Dissections were carried out at 4°C in ice-cold HBSS lacking MgCl2 and CaCl2 (HBSS−/−; Gibco, cat. no. 14180046). The dissected cortical tissue was subsequently transferred to a 15 mL tube containing 5 mL of ice-cold HBSS supplemented with MgCl2 and CaCl2 (HBSS+/+; Gibco, cat. no. 14025050). After centrifugation at 350 × g for 1 min at 4°C, the supernatant was aspirated and the tissue was resuspended in 10 mL of dissociation buffer comprising 8.6 mL HBSS+/+, 0.4 mL of 0.5 mg/mL DNase I (Roche, cat. no. 10104159001), and 1.0 mL of 2.5% trypsin (Gibco, cat. no. 15090046). The tissue was incubated at 37°C for 20 min, with gentle agitation every 5 min. After 10 min of incubation, mechanical dissociation was initiated by gently pipetting the tissue with a P1000 tip with the end cut off. At the end of the 20 min incubation, the tissue was further mechanically dissociated at the bottom of the tube by pipetting approximately 20 times with an uncut P1000 tip, until the suspension became cloudy and most visible tissue fragments had dispersed. The dissociation was stopped by centrifuging the single cell suspension at 350g x 1min and resuspending in ice cold PBS twice. The cell suspension was passed through 35 µm strainer caps into round-bottom tubes and maintained on ice until sorting on a BD FACS Aria Fusion flow cytometer equipped with a 100 µm nozzle. Cells were sorted into a 384-well plate, pre-loaded with heavy SILAC spike-in as prepared above.

#### Single-cell proteomics data analysis

Data pre-processing was performed according to the Methods section of Welter and Mutschler et al. 2026 ^48^, under “Data analysis”, following the subsections “Raw data processing”, “Processing of DIA-NN output” and “Technical noise”. Cells that passed the quality filters described in the original study were retained for further analysis. Additionally, cells with fewer than 1500 proteins and less than 650 light-channel precursors were excluded by q-value filtering. Technical noise was simulated for all proteins individually, and only intensities (medium-heavy or heavy) where the simulated technical CV^2^ was smaller than the total CV^2^ were retained. Cell size normalization was performed similar to what is described in the subchapter “Cell Cycle Models” of the original study ^48^. Instead of relying on FACS data, summed intensities per cell were used as estimator of cell size and a linear regression between total protein intensity and cell size was performed. Normalized protein intensities were calculated based on the residuals of the regression. Proteins detected in fewer than five cells and cells with less than 100 proteins were removed. M/L ratios were calculated from the medium-heavy and light-labeled intensities.

NA-aware distances between cells were estimated with dist_approx from proDA. The mean distance matrix was symmetrized and subjected to classical multidimensional scaling (MDS) using up to 20 dimensions The MDS coordinates were used to construct nearest-neighbour and shared-nearest-neighbour graphs in Seurat. Clustering resolution was 0.92. UMAP was calculated from the same MDS dimensions. Random seeds were fixed for clustering and UMAP. Cluster-associated proteins were identified separately for each assay using one-versus-rest proDA models on the original log10 matrix with missing values retained. P-values were adjusted by the Benjamini–Hochberg procedure.

## Acknowledgements

We thank the Technical Workshop of the Max Planck Institute for Molecular Genetics for support in designing and fabricating custom parts for the embryo roller culture unit. We are grateful for fruitful discussions with the Teresa Rayon lab during this project, and Aydan Bulut-Karslioglu, Florian Heyd, and Helene Kretzmer on M.C.-I’s thesis advisory committee.

M.L.K. was funded by the Minna-James-Heineman and Minerva Foundations, and the Deutsche Forschungsgemeinschaft (DFG) grant SPP2502 (563470554). E.O. was funded by the International Max Planck Research School for Biology and Computation, and both M.C.-I. and E.O. by grants from the German Academic Exchange Service (DAAD). D.V. and F.C. acknowledge funding support by the Federal Ministry of Education and Research (BMBF), as part of the National Research Initiatives for Mass Spectrometry in Systems Medicine (grant agreement No. 161L0222), funding from the European Research Council (ERC) under the European Union’s Horizon 2020 research and innovation program (grant agreement No. 101115681) and by the Initiative and Networking Fund of the Helmholtz Association within the framework of the Transfer Campaign (Helmholtz Co-Creation Projects). E.F.F. was supported by DFG grants SFB 1286 C12 and FO 1342/1-5, and NIH grant 1R21AG085062.

## Author contributions

M.L.K. designed and initiated the study. M.L.K. supervised the study with support from M.S., F.C., E.F.F., and K.I. The embryo culture and SILAC labeling method (MEMBRYO) was developed and performed by E.O. Computational analysis was performed by M.C.-I., with support from E.F.F., R.K., F.M., D.S.V., and A.S.W. Amino acid metabolomics were measured by B.L.-M. and D.M. Bulk proteomics samples were measured by R.K., single-cell proteomics by F.M. and A.S.W., and spatial proteomics by D.S.V., E.O., M.C.-I. and K.I. Data were interpreted by M.C.-I. and M.L.K. Manuscript figures and text were composed by M.C.-I. and M.L.K., with valuable editing and input from all authors.

## Materials and correspondence

Requests for materials and correspondence should be addressed to Matthew L. Kraushar.

---

**Extended Data Fig. 1.**
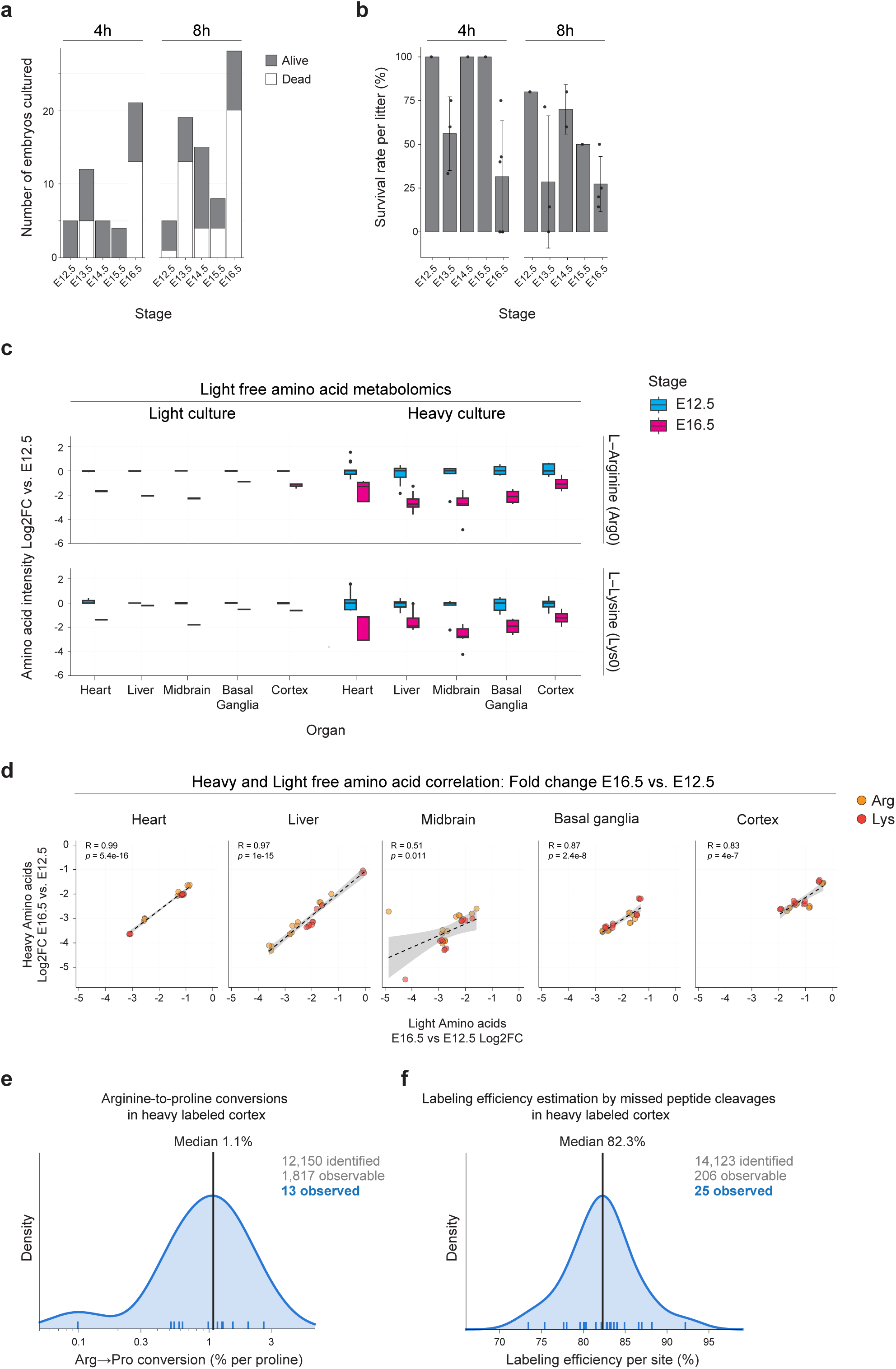
Embryonic viability and heavy free amino acid availability with MEMBRYO. **a**, Number of embryos alive or dead in this study after 4 or 8 hours of culture with heavy SILAC amino acids, at stages from E12.5 to E16.5. **b**, Survival rate per litter corresponding to (a). Each point represents one litter. **c**, The change in light arginine and lysine levels measured between E16.5 and E12.5 by metabolomics in five tissues, in both light (control) and heavy labeling cultures. Notably, light free amino acid levels decrease from E12.5 to E16.5 in all tissues and culture conditions tested. These data suggest a global change in the availability or metabolism of amino acids occurs during embryonic growth, which is supported by a previous study that measured decreasing histidine in the developing brain ^11^. **d**, Correlation in the levels of labeled heavy (y-axis) and pre-existing light (x-axis) arginine and lysine in tissues from embryos undergoing heavy culture. Each point represents the fold change from E12.5 to E16.5, and is a biological or a technical metabolomics replicate. Black dotted lines are the linear regression fits shown together with Pearson correlation coefficients (R) and associated p-values. **e**, Estimation of arginine-to-proline conversion rates in heavy SILAC labeled cortical tissue, combined data for each stage from E12.5-E15.5. **f**, Estimation of labeling efficiency based on missed peptide cleavages in cortical tissue, combined data for each stage from E12.5-E15.5. For **e-f**, “identified” denotes all peptides identified in the DDA dataset; “observable” denotes peptides meeting the sequence/composition criteria required for the respective analysis; “observed” denotes peptides for which both peptide forms needed to calculate the ratio were actually quantified.” E16.5 did not yield sufficient peptide coverage for estimation.

**Extended Data Fig. 2.**
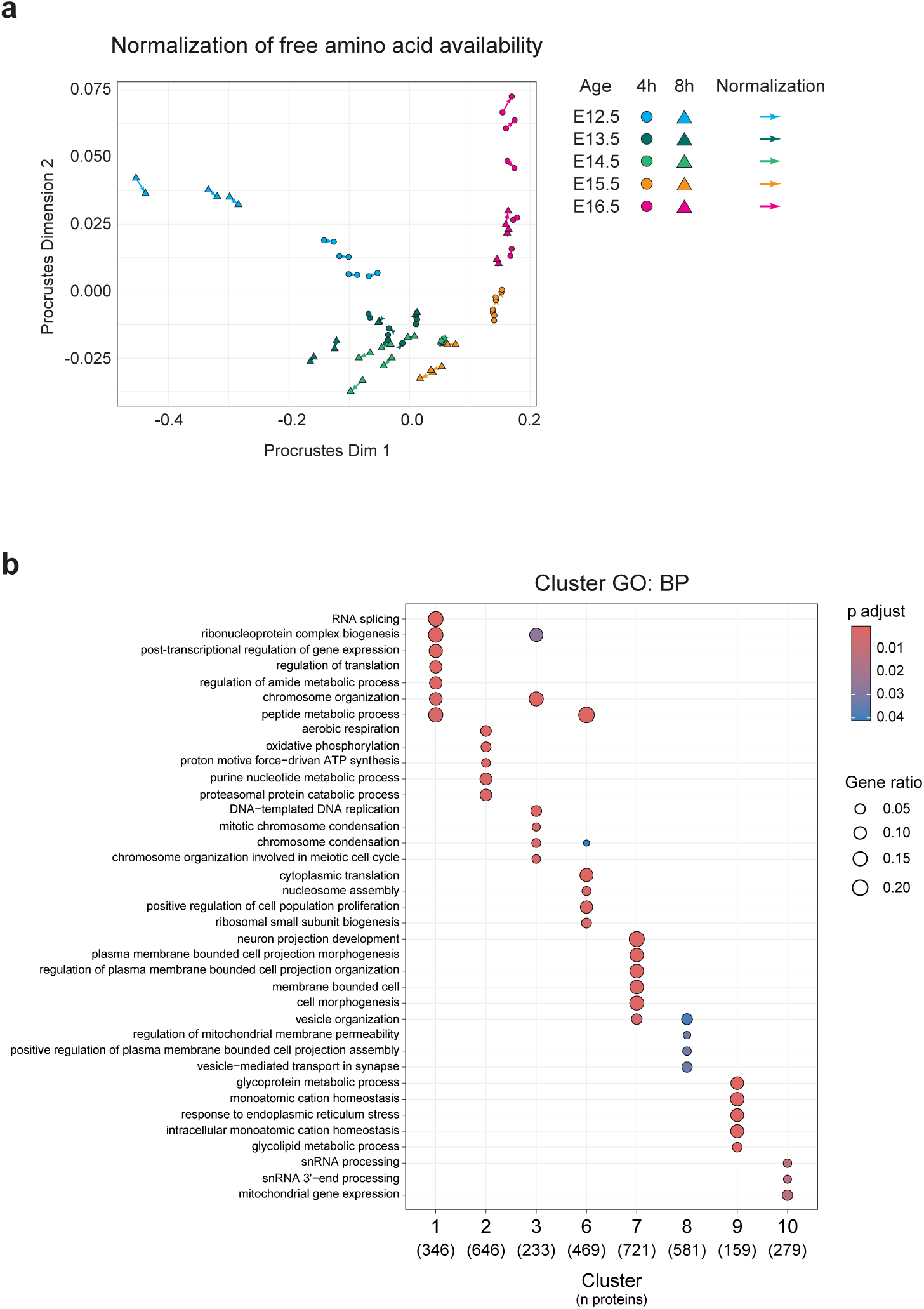
Evaluation of free amino acid normalization on heavy proteome intensity and pathways undergoing translation regulation. **a**, Procrustes superimposition of PCA configurations computed from heavy-to-total (H/T) ratios before and after free amino acid normalization, corresponding to Fig. 2b. The PCA configuration derived from the unnormalized H/T ratios was used as the reference, and the configuration derived from free amino acid-normalized H/T ratios was rotated and scaled to maximize correspondence. Each point represents one sample, and arrows connect the positions of matched samples before and after normalization. The minimal displacement of samples indicates near-perfect agreement between the two configurations (Procrustes correlation = 0.997; sum of squared residuals = 0.006). Statistical significance of the correspondence was assessed using a PROTEST permutation test with 999 permutations (P = 0.001). Colors indicate embryonic age, and shapes indicate embryo culture duration. **b**, Analysis of genes with distinct patterns of translation regulation throughout cortical development, measured by Ribo-seq and SILAC proteomics, and clustered by their trajectories over time in both assays (Fig. 2f). Shown is the enrichment of genes in each of the 10 identified clusters by gene ontology (GO) analysis of biological process (BP).

**Extended Data Fig. 3.**
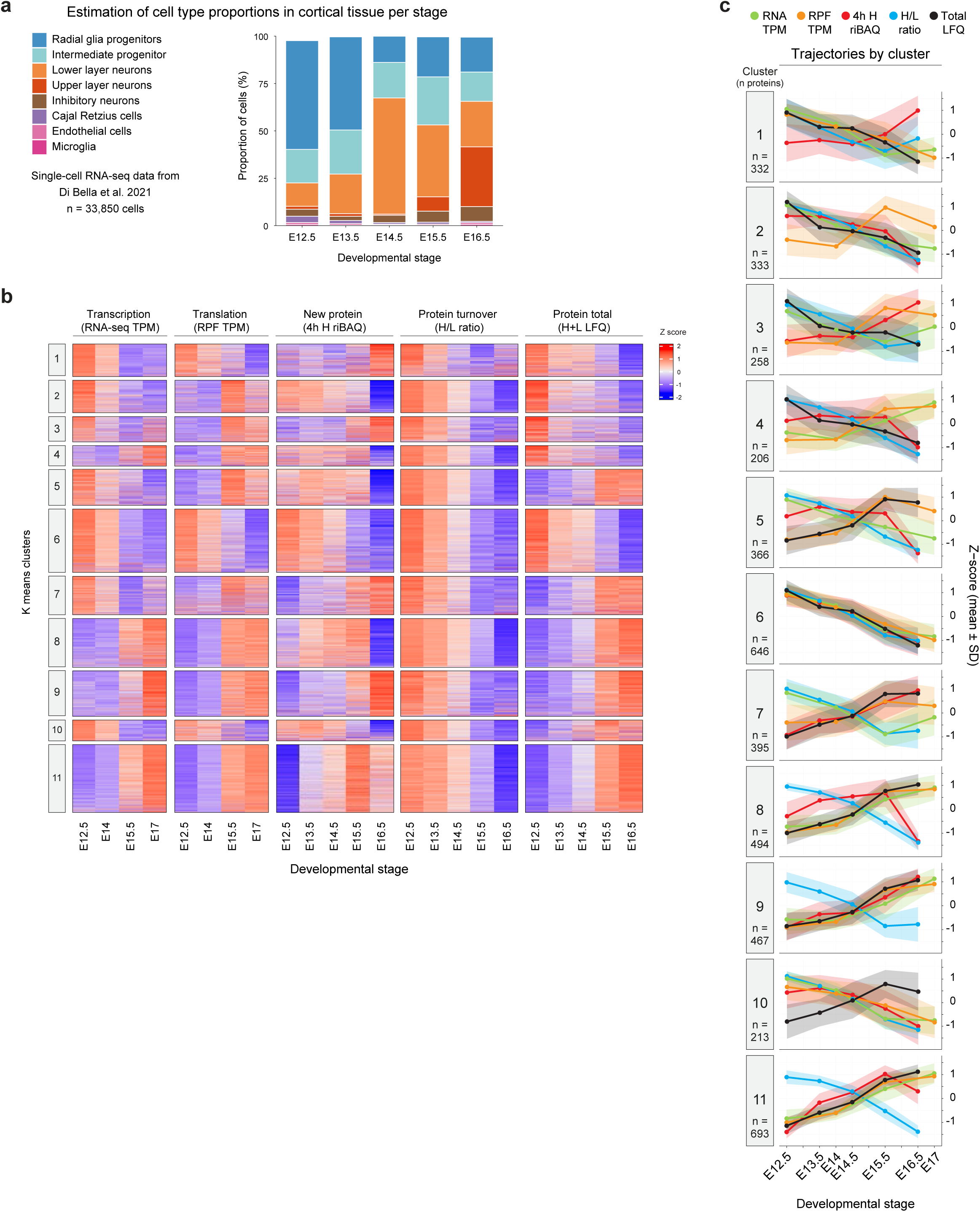
Changing cell types and patterns of transcriptional and post-transcriptional gene expression regulation in the developing cortex. **a**, Cell type composition across embryonic ages (E12.5-E16.5) estimated from scRNAseq data (Di Bella et al. 2021 ^26^; n = 33,850 cells). **b**, Heatmap of row-wise z-score normalized trajectories for RNA-seq TPM, Ribo-seq RPF TPM, SILAC riBAQ Heavy 4h, heavy-to-light protein intensity ratio, and LFQ total protein intensity in the cortex across embryonic ages. Rows represent individual proteins grouped by k-means cluster assignment (k = 11); rows are ordered within each cluster by distance to the cluster centroid. Cluster numbers are shown on the left. RNA-seq and Ribo-seq RPF TPM are derived from previously published data ^16^. **c**, Mean ± SD z-score trajectories per cluster as shown in (b), for the five molecular modalities measured across embryonic ages: RNA abundance as a proxy for transcription (RNAseq TPM, green), ribosome-protected fragment density as a proxy for translation (Ribo-seq RPF TPM), newly synthesized protein relative abundance (riBAQ Heavy 4h, red), heavy-to-light ratio as a proxy for turnover (H/L, blue), and total protein abundance (LFQ Total, black). Note that RNA-seq and Ribo-seq data span E12.5-E17 (E12.5, E14, E15.5, E17) ^16^, while SILAC data span E12.5-E16.5 (E12.5, E13.5, E14.5, E15.5, E16.5); both are plotted on the embryonic stage axis with correct intervals.

**Extended Data Fig. 4.**
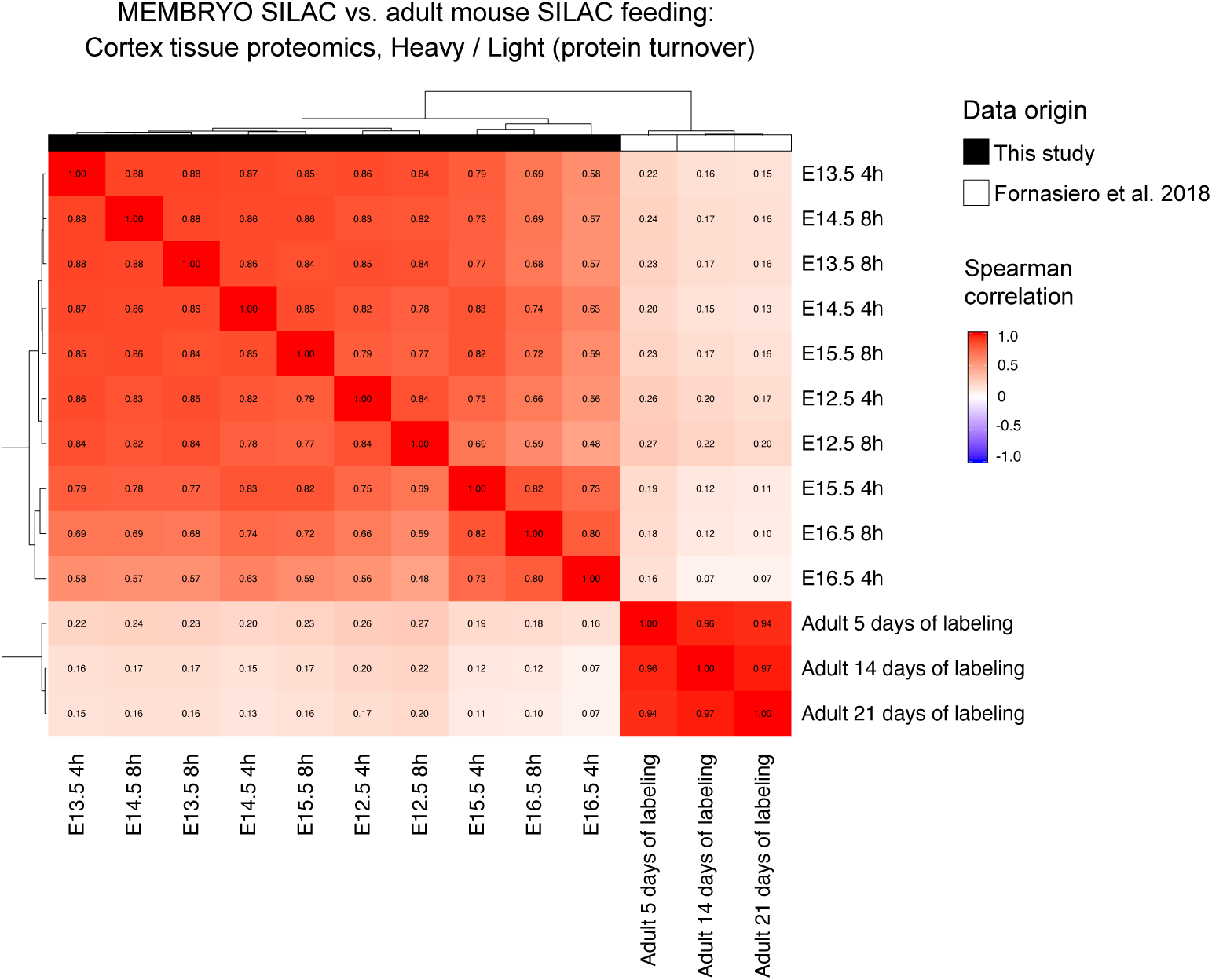
Per-protein turnover in the mouse embryonic and adult cortex. Spearman correlation of the heavy-to-light ratio (H/L) as a proxy for protein turnover, derived from the embryonic mouse cortex in this study (E12.5 to E16.5) vs. adult mouse cortex in Fornasiero et al, 2018 ^17^. The adult mouse cortex proteome was labeled by feeding mice with pellets containing isotopic amino acids for 5, 14 or 21 days.

